# A bipartite, mutation-tolerant NLS regulates interaction of ΔNp63α with importin alpha, nuclear transport and transcriptional activity

**DOI:** 10.1101/2025.06.12.659319

**Authors:** Anna Demarinis, Sara Esmaeili, Crystall M. D. Swarbrick, Simone Ranno, Silvia Pavan, Jade K. Forwood, Brian P. McSharry, Martin Pal, Enzo Di Iorio, Gualtiero Alvisi

**Affiliations:** Department, of Molecular Medicine, University of Padova, Padova, 35121; School of Dentistry and Medical Sciences, Faculty of Science and Health, Charles Sturt University, Wagga Wagga, NSW, Australia; Gulbali Institute, Charles Sturt University, Wagga Wagga, NSW, Australia

## Abstract

ΔNp63α is a master regulator of epithelial development, driving the expansion of progenitor cells in stratified epithelia. Mutations in ΔNp63α are linked to squamous cell carcinomas (SCCs) and basal cell carcinomas (BCCs), as well as to ectodermal dysplasia syndromes such as ectrodactyly-ectodermal dysplasia-clefting (EEC) and ankyloblepharon-ectodermal dysplasia-clefting (AEC). Although ΔNp63α functions as a nuclear transcription factor, the mechanisms underlying its nuclear import remain incompletely understood. By combining imaging, biochemical, structural and functional assays, we have thoroughly characterized ΔNp63α nuclear import, as mediated by the importin (IMP) α/β1 heterodimer. We also show here that ΔNp63α has evolved a peculiar strategy to ensure mutation tolerant nuclear localization, which is essential for DNA binding and transcriptional regulation. Despite a canonical bipartite NLS formed by two stretches of basic amino acids was identified between the DNA binding and oligomerization domains, each of them in sufficient to bind both IMPα binding sites upon homodimerization. Therefore, in contrast to most known bipartite NLSs, only simultaneous substitution of both basic stretches of amino acids ablated nuclear localization, interaction with IMPα, and decreased transcriptional activity. Since several ΔNp63α isoforms which lack the N-terminal basic stretch of amino acids have been described, and a number of mutations in the ΔNp63α NLS region have been identified in the Genome Aggregation Database, ΔNp63α has specifically evolved to tolerate mutations in its NLS without significantly compromising its ability to localize in the nucleus.

**GRAPHICAL ABSTRACT:** 

## INTRODUCTION

The p53 transcription factor family consists of the tumor suppressor p53 and the orthologous proteins, p63 and p73 (1, 2). Unlike p53, p63 plays an important role in the development, differentiation, and morphogenesis of various epithelial tissues (3, 4). As a result of a complex transcriptional regulation process that involves the use of multiple promoters and alternative splicing events, several p63 isoforms are expressed in epithelial tissues and their appendixes, each endowed with specific and sometimes contrasting functions (5). All p63 isoforms share a DNA binding domain (DBD) and an oligomerization domain (OD), along with distinct domains at both the N- and C-terminus (6). Based on the use of alternative promoters, the p63 gene can give rise to nger TA isoforms, or shorter ΔN isoforms. TA isoforms contain an N-terminal transactivation domain (TAD), whose sequences are conserved among p53 family proteins to transactivate target gene expression, while ΔNp63 proteins lack the entire TA domain and instead have ΔN-specific sequences of only 14 amino acid residues (7). As a result of a complex alternative splicing process, TA and ΔNp63 isoforms can be further classified as α, β, γ, δ, or ε, based on the presence of different additional domains at the C-terminus, which include another TAD (named TAD2), a sterile alpha motif (SAM), and transactivation inhibition domain (TID; (8)). Furthermore, TAp63 and ΔNp63 isoforms are present during distinct stages of embryonic development. During epidermal development, while the TAp63 variants are expressed in the uncommitted ectoderm surface, the ΔNp63 isoforms are transcribed only after the surface ectoderm has been committed towards a stratification program. Mice lacking ΔNp63 exhibit single-layered surface epithelium instead of a fully stratified epidermis, displaying a dramatic loss of keratinocytes (4, 9). ΔNp63α is the prominent isoform in the adult epidermis, where it contributes to self-renewal ability or stemness (10), and is frequently overexpressed in a variety of squamous cell (SCC) and basal cell carcinomas (BCCs; (11)). Based on their structural characteristics and their ability to inhibit p53 and TA-p63, ΔNp63 proteins were originally defined as dominant-negative type suppressors against the p53 family (9), even though more recent data demonstrated their role as transcriptional activators of specific genes (12). Indeed, the different p63 isoforms are endowed with the ability to differently regulate gene expression (13). Accordingly, ΔNp63α is known to either repress or stimulate transcription from different promoters, inhibiting p53-mediated transcriptional activation (14) and stimulating transcription from the K14 promoter (13). In both cases, transcriptional regulation depends on the ability to bind specific dsDNA elements upon tetramerization (15, 16). Furthermore, mutations in the DBD have been linked to several genetic diseases including ectrodactyly-ectodermal dysplasia-clefting syndrome (EEC) (17).

To function as gene regulators, transcription factors such as p53 family members must localize to the nucleus by traversing the nuclear envelope (NE) (18). Passage across the NE occurs through large macromolecular complexes called nuclear pore complexes (NPCs), composed of multiple copies of ∼30 different proteins known as nucleoporins (Nups), which form aqueous channels allowing passive diffusion of ions and proteins with a molecular weight (MW) below 40-70 Kd (19, 20). Larger proteins, or those that need to quickly and strongly accumulate in the cell nucleus such as transcription factors are instead actively transported across the NPC by specific nuclear transport receptors (NTRs) belonging to the karyopherin superfamily (21). NTRs can be further divided into importins (IMPs) and exportins (EXPs), which transport cargoes from the cytoplasm to the nucleus and vice-versa, respectively. Usually, IMPβ1 or one of its 20 orthologues recognize specific nuclear localization (NLS) or nuclear export signals (NES) on cargo protein before mediating translocation across the NPC. Each of the NTRs has been reported to recognize specific groups of cargoes so that proteins with similar functions are believed to be transported by the same subset of NTRs (22, 23). The best-characterized NLSs are called “classical”, and are recognized by IMPα1 or one of its six orthologues, which function as adaptors bridging cargoes and IMPβ1 (24). Classical (c)NLSs are highly basic sequences and can be further divided into monopartite NLSs, formed by a single stretch of basic amino acids binding to IMPα major binding site (25), or bipartite NLSs formed by two stretches of basic amino acids, separated by a short linker of 8-20 amino acids, both essential for IMPα binding and nuclear transport (26–32). Since ΔNp63α monomers have an apparent MW of 75 kDa, well above the threshold for passive diffusion through NPCs (20), they need to functionally interact with NTRs to access the nucleus (33). Furthermore, p53 family members’ subcellular localization, levels, and transcriptional activity are finely regulated thanks to interaction with specific Nups and NTRs (33). Accordingly, both p53 and p73 bear bipartite NLSs and NESs, regulating their subcellular localization and transcriptional properties (31, 34–36). However, little is known regarding p63 nucleocytoplasmic transport, the NLSs involved, and the NTRs responsible for the process. A recent study described an NLS involved in the nuclear transport of ΔNp63α in an IMPβ1 and Nup62-dependent fashion (37). However, the inactivation of such NLS was not sufficient to prevent nuclear accumulation, precluding more detailed studies regarding the relationship between ΔNp63α function and its subcellular localization, and raising the question regarding the contribution of specific sequences to ΔNp63α nuclear import (37).

In this study, we have thoroughly characterized the nuclear import of ΔNp63α using quantitative confocal laser scanning microscopy (CLSM), demonstrating that it is dependent on the IMPα/β1 transport pathway. We also present a detailed atomic-level structural analysis of the ΔNp63α-NLS:IMPα complex, revealing that p63 residues 278-DGTKRPFRQNTHGIQMTSIKKRRSP-302 bind across the inner face of IMPa. Specifically, the N-terminal stretch, KRPFR, interacts at the minor binding site, while the C-terminal stretch, KKRR binds at the major site. Such NLS interacts with several IMPa isoforms, exhibiting a preference for IMPa1, 3, and 5 over IMPa2 and IMPa7, while not binding directly with IMPβ1. Substitution of key residues within either basic stretch of basic amino acids forming the p63 bipartite NLS strongly affected IMPa binding and nuclear accumulation when the newly identified NLS was studied in isolation, as shown for typical linear bipartite NLSs. However, the same substitutions only partially affected nuclear import of full-length ΔNp63α, and only simultaneous inactivation of both basic stretches of amino acids precluded nuclear accumulation. Further, by combining CLSM, bioluminescent resonant energy transfer (BRET), and AlphaFold modelling, we could demonstrate that ΔNp63α can self-interact in the cytoplasm and bind to IMPa as a dimer, whereby the high flexibility of the NLS region allows alternative binding modes. As a result, the same basic stretch of amino acids from different monomers is predicted to interact simultaneously with both IMPa major and minor binding sites. Since we show that nuclear localization is crucial for ΔNp63α DNA binding and transactivation properties, the evolution of such three-dimensional NLS ensures nuclear targeting of ΔNp63α even in the presence of rare mutations with the newly identified ΔNp63α NLSbip reported in the human population, further stressing the functional relevance of the NLS identified in this study.

## MATERIAL AND METHODS

### Bioinformatics

The primary sequence from ΔNp63α (UniProt code; Q9H3D4-2), was analyzed with the cNLS mapper software (38), and by visual inspection to identify putative NLSs. ΔNp63α isoforms containing the full NLSbip (DGT**KR**PF**R**QNTHGIQMTSI**KKRR**SPD) and its alternatively spliced variant lacking the N—terminal basic stretches of amino acids (DAF**R**QNTHGIQMTSI**KKRR**SPD) were retrieved searching the non-redundant protein database using the BLASTP algorithm (39). Structural models were predicted using the AlphaFold3 Server (40) and visualized using the PyMOL Molecular Graphics System (version 3.1.3.1; Schrödinger, LLC). Protein model interactions were analyzed using the PDBePISA (https://www.ebi.ac.uk/pdbe/pisa/) and PDBsum (https://www.ebi.ac.uk/thornton-srv/databases/pdbsum/) web tools (41).

### Plasmids

Bacterial expression plasmids mediating the expression of the truncated version of human IMPα1, IMPα3, IMPα5, and IMPα7 and mouse (m)IMPα2 lacking the autoinhibitory IMPβ1 binding (IBB) domain, and mIMPβ1 from a pET30a backbone were described previously (42, 43). Mammalian expression plasmids pcDNA3.1-NT-GFP-TOPO and pcDNA3.1-NT-GFP-TOPO-SV40-NLS, mediating the expression of GFP cycle3, or of a fusion protein between GFP cycle3 and Simian vacuolating virus (SV) 40 large tumor antigen NLS (PKKKRKV-132), respectively, were described previously (44). Plasmid pEPI-GFP-UL44, encoding a fusion protein between GFP and human cytomegalovirus (HCMV) DNA polymerase UL44, localizing to the nucleus via the IMPα/β1 dependent pathway, was also previously described (45). Plasmid mCherry-Bimax2, encoding a competitive inhibitor of the IMPα/β1 nuclear import pathway (46), was a generous gift from Yoshihiro Yoneda and Masahiro Oka (Osaka, Japan). Mammalian expression plasmids encoding GFP-p53, GFP-ΔNp63α, and substitution derivatives thereof, as well as GFP fused to ΔNp63α bipartite NLS, were generated by BioFab research (Rome, Italy), by cloning synthesized ORFs in the expression plasmid pEGFP-C1 (Clontech). pDNR207plasmids were generated by Gateway^TM^ BP recombination reactions, and Mammalian expression plasmids mediating expression of FLAG-, YFP-RFP- and Renilla luciferase (RLuc) fusion proteins were generated by Gateway^TM^ LR recombination reactions between pDNR207 plasmids and pDESTntYFP, pDESTntRFP, pDESTntRLuc (47) or pDESTntFLAG (48) destination vectors. Empty Vector pcDNA3 (#V79020, Thermo Fisher Scientific, Monza, Italy), used to normalize transfection amounts was purchased. Plasmid Tag-RFP-N, mediating the constitutive expression of a bright monomeric red fluorescent protein (49), was a generous gift from Roger Tsien (San Diego, USA). Plasmid 2xP53_RE::NLuc (Addgene #124533), whereby Nanoluficerase (NLuc) expression is under the control of two p53 responsive elements (RE), was described in (50), and generously gifted by Koen Venken (Ghent, Belgium). A list of all plasmids used in this study is available in Supplementary Table S1.

### Peptides

Synthetic peptides representing the predicted amino acid residues, with an N-terminal FITC tag, were synthesized by GenScript (Supplementary Table S2).

### Expression and purification of recombinant proteins

The overexpression of IMPs was carried out in *E. coli* pLysS BL21(DE3) as described previously (51) using the auto-induction method (52). After induction, the cultures were centrifuged at 6,000 rpm for 20 minutes at 4°C, and the resulting bacterial pellets were resuspended in HIS buffer A (pH 8), which consisted of 50 mM phosphate buffer, 300 mM NaCl, and 20 mM imidazole. To lyse the cells, two freeze-thaw cycles were performed, followed by the addition of lysozyme (1 mL of 20 mg/mL; #L6876, Sigma-Aldrich, St. Louis, MI, USA) and DNase (10 μL of 50 mg/mL; #DN25, Sigma-Aldrich, St. Louis, MI, USA), and incubation at room temperature for 1 hour. The supernatants containing soluble proteins were collected by centrifugation at 12,000 rpm for 30 minutes at 4°C. The extracts were then filtered through 0.45 μm low protein affinity filters and injected into a 5 mL HisTrap HP column (#71-5027-68 AF, GE Healthcare, Chicago, IL, USA) that had been pre-equilibrated with His buffer A, in an AKTA purifier FPLC system (GE Healthcare, USA). Followed by washes of 20 column volumes with His buffer A, the proteins of interest were eluted using a gradually increasing gradient of imidazole (ranging from 20 mM to 500 mM; #IM00260250, ChemSupply, Gillman, SA, Australia). The eluted protein fractions were combined and loaded onto a pre-equilibrated HiLoad 26/60 Superdex 200 column (#28-9920-17 AC, GE Healthcare, USA) in GST buffer A (50 mM Tris and 125 mM NaCl) for further purification using size-exclusion chromatography. The fractions corresponding to the eluted volumes at the respective protein sizes were collected, and the samples were concentrated using an Amicon MWCO 10 kDa filter (#UFC9010, Merck Millipore, Burlington, MA, USA). Before experimental use, the purity of the samples was assessed by SDS-PAGE on a 4–12% Bis-Tris Plus gel (#NW04120BOX, Thermo Fisher Scientific, Waltham, MA, USA).

### Fluorescence polarization (FP) assays

FITC-tagged peptides (Supplementary Table S2) at a concentration of 2 nM were incubated with IMPα1ΔIBB, mIMPα2ΔIBB, IMPα3ΔIBB, IMPα5ΔIBB, IMPα7ΔIBB and IMPβ1 in two-fold serial dilutions, starting from 20 μM, across 23 wells to a final volume of 200 μL per well in GST buffer A (50 mM Tris, 125 mM NaCl), with the last well containing only GST buffer A, as described previously (53, 54). FP measurements were conducted using a CLARIOstar Plus plate reader (BMG Labtech, Germany) in 96-well black Fluotrac microplates (#10701634, Greiner Bio-One, Austria). Assays were carried out in three independent experiments. Data were analyzed with GraphPad Prism (Prism 9, Version 9.3.1), and a binding curve was fitted to determine the dissociation constant (K_d_) and maximum binding (B_max_). A summary of the FP assay results is presented in Supplementary Table S3.

### Crystallization, data collection, and structure determination

The hanging drop vapor diffusion method was employed to resolve the IMPα2ΔIBB:p63 NLS protein complex. IMPα2ΔIBB was first crystallized, with each hanging drop having a total volume of 3 μL over a well of 300 μL precipitant solution containing 650 mM sodium citrate, 100 mM HEPES (pH 6.5), and 10 mM DTT at a temperature of 23°C. Crystals were formed after 3 days of incubation. Formed crystals were then soaked with peptide to obtain a 1:1 complex, collected and cryoprotected in a precipitant solution containing 20% glycerol, before being rapidly frozen in liquid nitrogen. X-ray diffraction data were obtained at the Australian Synchrotron using the MX2 macromolecular beam line, utilizing an Eiger 16M detector (55). The data obtained were subjected to indexing and integration using XDS (56). Subsequent steps including merging, space group assignment, scaling, and Rfree calculations were carried out using AIMLESS (57) within CCP4 (58). Phasing was performed using molecular replacement in Phaser (59) and PDB code 6BW1 was used as the search model for IMPα2. Model building and refinement were performed in iterative rounds using COOT (60), and Phenix (61). The finalized model was subjected to validation and subsequently deposited to the Protein Data Bank (PDB) with an assigned accession number 9N54; refinement statics are detailed in Supplementary Table S4. The binding interactions were calculated using PDBePISA (41).

### Cells

The non-small-cell lung carcinoma H1299 cell line (#CRL-5803, ATCC), human embryonic kidney (HEK)293T (#CRL-3216, ATCC) and HEK293A (#R70507, Thermo Fisher Scientific, Monza, Italy) cells were maintained in Dulbecco’s modified Eagle’s medium (DMEM) supplemented with 10% (v/v) fetal bovine serum (FBS), 50 U/ml penicillin, 50 U/ml streptomycin, and 2 mM L-glutamine in a humidified incubator at 37°C in the presence of 5% CO_2_ and passaged when reached confluence as described in (62).

### Transfections

HEK293A cells were seeded in a 24-well plate onto glass coverslips (5×10^4^ cells/well). HEK293T (1×10^5^ cells/well) and H1299 (6×10^4^ cells/well) were seeded in a 24-well plate. The next day, cells were transfected with appropriate amounts of expression constructs (range 5-250 ng), using Lipofectamine 2000 (#11668019, Thermo Fisher Scientific, Monza, Italy), following the manufacturer’s recommendations and further incubated at 37 °C and 5% CO_2_ in complete DMEM as described (63), until being processed for CLSM, BRET or transactivation assays.

### CLSM and image analysis

Transfected cells were incubated for 24 hours to allow the expression of spontaneously fluorescent proteins. Afterward, the cells were treated with DRAQ5 (#62251, Thermo Fisher Scientific, Monza, Italy) at a dilution of 1:5,000 in DMEM without phenol red for 30 minutes (64). Following incubation, the cells were washed twice with PHEM 1x solution (60 mM PIPES, 25 mM HEPES, 10 mM EGTA, and 4 mM MgSO_4_) and fixed with 4% paraformaldehyde (v/v) for 10 minutes at room temperature (RT). After three washes with PBS 1x, coverslips were mounted on glass slides using Fluoromount G (#00-4958-02, Thermo Fisher Scientific, Monza, Italy). The subcellular localization of fusion proteins was examined using a Nikon A1 confocal laser scanning microscope (Nikon, Tokyo, Japan) equipped with a 60x oil immersion objective, following the established protocol outlined previously (65, 66). To determine the levels of nuclear accumulation of the proteins of interest, the FiJi public domain software (https://doi.org/10.1038/nmeth.2019) was utilized, and single-cell measurements were taken for nuclear (Fn) and cytoplasmic (Fc) fluorescence. DRAQ5 was used to define nuclear masks, while a small area close to DRAQ5 was used to define a cytosolic mask as previously (67). The fluorescence attributed to autofluorescence/background (Fb) was subtracted from the measurements, to calculate the Fn/c ratio according to the formulas Fn/c = (Fn-Fb)/(Fc-Fb). Cells with oversaturated signals were excluded from the analysis. Statistical analysis was performed using GraphPad Prism 9 software (GraphPad, San Diego, CA, USA) applying either Student’s t-test, one-way ANOVA, or two-way ANOVA, as appropriate.

### Bioluminescent resonant energy transfer (BRET) assays

BRET experiments were performed in HEK293T cells as described in (62, 63, 68). For each construct, the donor (RLuc)-expressing plasmid was transfected both in the absence and the presence of the relative acceptor (YFP)-expressing plasmid to allow calculation of background BRET signal. At 24 h post-transfection, the culture medium was removed from wells, and cells were gently washed with 1 mL of PBS before being resuspended in 290 μL of PBS. Cell resuspensions (90 μL) were then transferred to a 96-well black flat-bottom polystyrene TC-treated microplate (#3916, Corning, Corning, NY, USA) in triplicate, and signals were acquired using a Varioskan^TM^ Lux multimode microplate reader compatible with BRET measurements (Thermo Fisher Scientific, Monza, Italy). Fluorescence signals (YFPnet) relative to YFP fluorescence emission were acquired using a fluorometric excitation filter (band pass 485 ± 14 nm) and a fluorometric emission filter (band pass 535 ± 25 nm). Luminometric readings were performed at 5, 15, 30, 45, and 60 min after the addition of h-Coelenterazine (PJK Biotech, Kleinblittersdorf, Germany; 5 μM final concentration in PBS). Data were acquired for 1 s/well, using a luminometric 530 ± 30 nm emission filter (YFP signal) and a luminometric 480 ± 20 nm emission filter (RLuc signal). Before reading, the plate was shaken for 1 s at normal speed and with a double orbit. After background subtraction using values relative to mock-transfected cells, the data obtained were used to calculate the BRET signal, defined as the ratio between the YFP and RLuc signals calculated for a specific BRET pair, according to the formula:

BRET signal = (YFP emission)/(RLuc emission).

Similarly, the BRET ratio, defined as the difference between the BRET value relative to a BRET pair and the BRET value relative to the BRET donor alone, was calculated according to the formula:

BRET ratio = [(YFP emission)/(RLuc emission) BRET pair] - [(YFP emission)/(RLuc emission) BRET donor].

### p53-mediated transcription inhibition assays

H1299 cells transfected with fixed amounts (40ng/well) of Tag-RFP-N and 2xP53-RE-NLuc reporter plasmids in the absence or the presence of pDESTnt-YFP53. When required, different amounts pDESTnt-FLAG-ΔNp63α or pDESTnt-FLAG-ΔNp63α;mNLSbip were also added (range 0-360 ng). Total transfection amounts were normalized to 500 ng with pCDNA3. Twenty-four hours post transfections, cells were washed with ice cold PBS1x, and 290 μl of passive lysis buffer (# E1941, Promega Italia, Milan, Italy) was added to each well. Cells were frozen by incubating 1 hour at -80°C, thawed on ice, and 100 μl transferred to a 96-well black flat-bottom polystyrene TC-treated microplate (#3916, Corning, Corning, NY, USA), and signals were acquired using a Varioskan^TM^ Lux multimode microplate reader compatible with fluorescence and luminometric measurements (Thermo Fisher Scientific, Monza, Italy). YFP and RFP Fluorescence signals were acquired by excitation at 488 and 530±12 nm, and acquisition at 555 and 583±12 nm, respectively. Luminometric readings were performed at 30 min after the addition of 100 μl/well of Luciferase Assay Buffer containing 2 μl Luciferase Assay Substrate from the Nano-Glo® Luciferase Assay System (#N1110; Promega Italia, Milan, Italy). Data were acquired for 1 s/well, with Automatic sensitivity (NLuc signal). Before reading, the plate was shaken for 1 s at normal speed and with a double orbit. After background subtraction using values relative to mock-transfected cells, the data obtained were used to calculate the transfection normalized p53 levels (YFP fluorescence/RFP fluorescence), as well as the transfection normalized p53 activity (NLuc luminescence/RFP fluorescence) and the intrinsic p53 activity (NLuc luminescence/YFP fluorescence). Data was further normalized to values obtained for cells transfected in the absence of inhibitors.

#### Interpretation of Sequence Variants

Variants present in the NLS sequence of the TP63 gene and their frequency were retrieved from the Genome Aggregation Database (gnomAD) vs. 4.1 a population dataset of over one million unrelated individuals, using the minor allele frequency (MAF) cut-off of 0.001 (69). The National Center for Biotechnology Information (NCBI) ClinVar (https://www.ncbi.nlm.nih.gov/clinvar), was used to aggregate information on genomic variation and its relationship to human health, particularly their clinical outcomes. The clinical impact of sequence variants was evaluated using the guidelines developed by the American College of Medical Genetics and Genomics (70).

## RESULTS

### ΔNp63α nuclear import is dependent on IMPα/β1

Although ΔNp63α nuclear import was recently proposed to depend on an NLS located at residues 295-304 (SIKKRRSPDD), and IMPβ1 (71), its NLS perfectly matches the consensus for binding to IMPα/β1 (K-K/R-X-K/R, see (66)). Therefore, we hypothesized the protein could be imported into the nucleus by IMPα/β1. To test this hypothesis, we expressed GFP-ΔNp63α in the absence or presence of the well-known IMPα/β1 inhibitor, Bimax2, and quantitatively analyzed its levels of nuclear accumulation at the single cell level by CLSM. As controls, we also expressed GFP alone, which lacks specific targeting signals and can freely diffuse across the NPC, as well as fusion proteins between GFP and HCMV DNA polymerase processivity factor UL44, which is transported into the nucleus by the IMPα/β1 heterodimer (72), and GFP with H1E, which is transported into the nucleus by multiple IMPs (73) (Figure 1A and Supplementary Figure S1). As expected, while GFP was evenly distributed throughout the cell, all GFP-fusion proteins exhibited strong nuclear localization when expressed in the absence of mCherry-Bimax2 (Figure 1B). Importantly, co-expression with mcherry-Bimax2 strongly impaired nuclear accumulation of GFP-UL44, but had no effect on GFP-H1E (Figure 1B-D). Co-expression with mcherry-Bimax2 almost completely abrogated GFP-ΔNp63α nuclear localization (Fn/c decreased from 40.2 to 0.9, see Figure 1C). Importantly, in the presence of mCherry-Bimax2 GFP-ΔNp63α failed to accumulate in the nucleus in 93% of co-transfected cells (Figure 1D). These findings indicate that ΔNp63α nuclear import is highly dependent on IMPα/β1.

**Figure 1.**
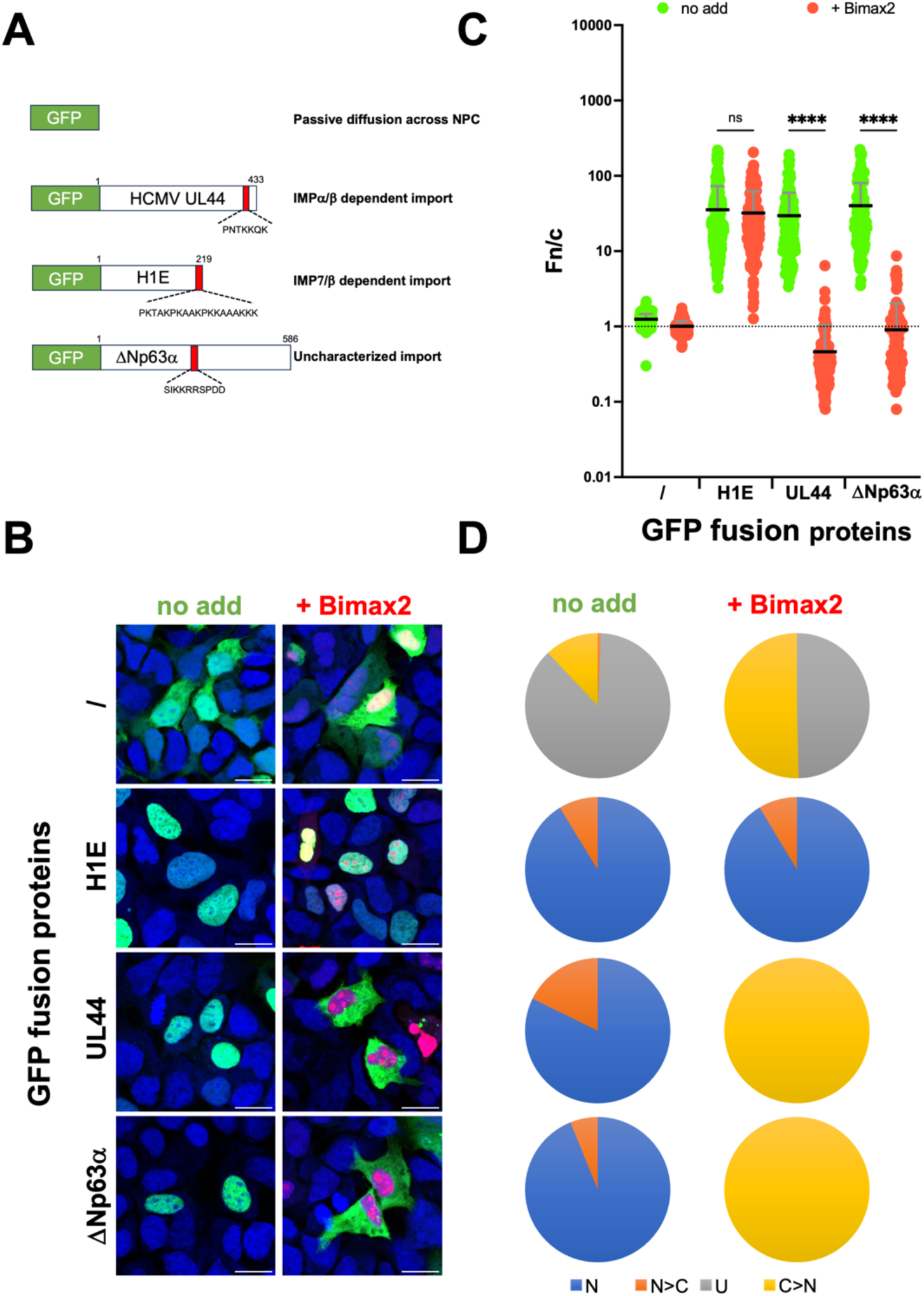
ΔNp63α is transported into the nucleus by the IMPα/β1 heterodimer. (A) HEK293A were transfected to express the indicated GFP fusion proteins, in the presence or the absence of the well-characterized IMPα/β inhibitor Bimax2. (B) Twenty-four hours later cells were incubated with DRAQ5 to stain cell nuclei, fixed, and processed for single-cell quantitative CLSM analysis as described in the Materials and Methods section. Merge images of the 633 (DRAQ5, nuclei), 561 (mcherry, Bimax2), and 488 nm (GFP, fusion protein) laser lines are shown. Individual cells expressing the indicated proteins were analyzed to calculate the Fn/c value relative to GFP fusion proteins. (C) The Fn/c values relative to the indicated fusion proteins are shown as individual measurements from single cells (circles), means (black horizontal lines) and standard deviation of the mean (grey vertical lines), relative to pooled data from three independent experiments, including the results of the Welch and Brown-Forsythe one-way ANOVA for significance between the indicated proteins. ****: p < 0.0001; ns: not significant. (D) Pie charts representing the percentage of cells displaying the indicated subcellular localizations. N: nuclear, Fn/c ≥ 10; N>C: nuclear more than cytosolic, 2 ≤ Fn/c < 10; U: ubiquitous, 1 ≤ Fn/c < 2; C>N: more cytosolic than nuclear, Fn/c < 1.

### ΔNp63α residues 278-302 mediate nuclear localization through two basic amino acids stretches

Since the nuclear accumulation of GFP-ΔNp63α was strongly impaired by the IMPα/β1 inhibitor Bimax2, but inactivation of its previously described NLS only caused a very minor impairment of nuclear import (71), we hypothesized that additional NLSs may contribute to IMPα/β1-dependent p63 nuclear import. Analysis of ΔNp63α primary sequence with the software cNLS mapper revealed a potential bipartite cNLS that partially overlaps with the previously identified NLS (278-DGTKRPFRQNTHGIQMTSIKKRRSP-302, see Figure 2A). To assess the functionality of the newly identified bipartite cNLS (NLSbip), we tested its ability to mediate GFP nuclear localization using quantitative CLSM. We also examined the impact of substituting either the N-terminal (mNLSn: 278-DGTaaPFRQNTHGIQMTSIKKRRSP-302), C-terminal (mNLSc: 278-DGTKRPFRQNTHGIQMTSIaaaaSP-302) or both (mNLSbip: 278-DGTaaPFaQNTHGIQMTSIaaaaSP-302) stretches of basic residues on nuclear targeting (Figure 2B). Importantly, the ΔNp63α NLSbip enhanced the nuclear targeting of GFP, which would otherwise be evenly distributed between the nucleus and the cytoplasm (Figure 2C and Supplementary Figure S2). Indeed, GFP-NLSbip accumulated into the nucleus of 75% of cells (Figure 2D), with a mean Fn/c of 2.6 (Figure 2E), significantly higher than that of GFP alone (Fn/c of 1.2, see Figure 2E). Substitution of the N-terminal stretch of basic amino acids (mNLSn; K281A/R282A), significantly reduced nuclear accumulation, resulting in a decrease of the Fn/c to 1.4, whereas substitution of the C-terminal stretch of basic amino acids (mNLSc; K297A/K298A/R299A/R300A) completely ablated NLS function (Fn/c of 1.2, see Figure 2E).

**Figure 2.**
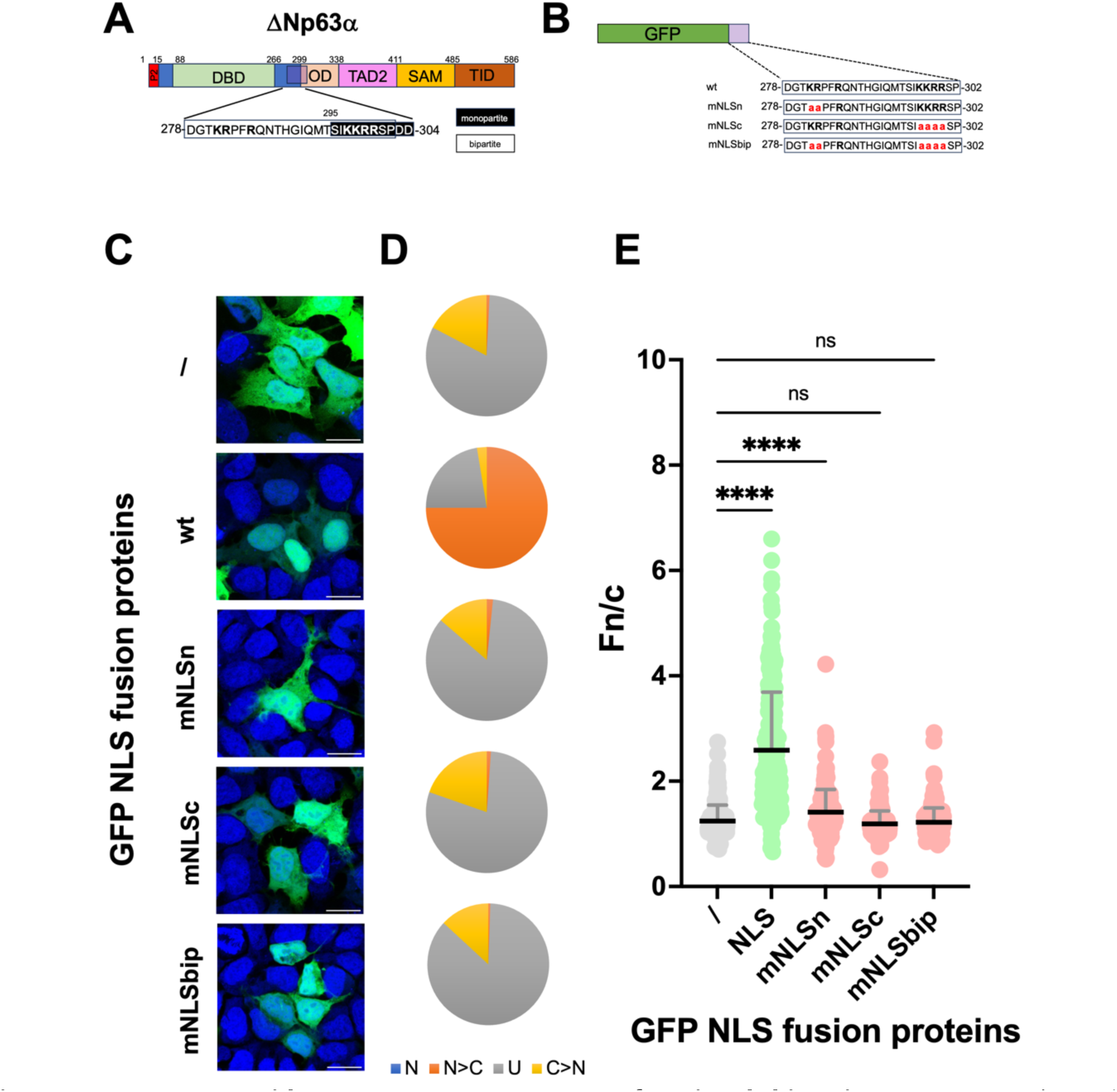
ΔNp63α residues 278-302 represent a functional bipartite NLS. (A) Schematic representation of ΔNap63 functional domains and NLSs. The sequence of a previously described monopartite NLS is boxed with a black background, whereas the putative bipartite NLS identified here is boxed with a white background. The single-letter amino acid code is used, with basic residues in boldface. (B) HEK293A cells were transfected to express the indicated GFP-NLS fusion proteins. The sequence of ΔNap63 NLSbip is shown as capital letters, with basic amino acids in boldface and substituted amino acids in red lowercase (C) Twenty-four hours later cells were incubated with DRAQ5 to stain cell nuclei, fixed, and processed for single cell quantitative CLSM analysis as described in the Materials and Methods section. Merged images of the 633 (DRAQ5, nuclei), 561 (mcherry, Bimax2), and 488 nm (GFP-NLS) laser lines are shown. Individual cells expressing the indicated proteins were analyzed to calculate the Fn/c value relative to GFP fusion proteins (D) Pie charts representing the percentage of cells relative to each indicated fusion protein displaying the indicated subcellular localizations are shown. N: nuclear, Fn/c ≥ 10; N>C: nuclear more than cytosolic, 2 ≤ Fn/c < 10; U: ubiquitous, 1 ≤ Fn/c < 2; C>N: more cytosolic than nuclear, Fn/c < 1. (E) The Fn/c values relative to the indicated fusion proteins are shown as individual measurements from single cells (circles), means (black horizontal lines) and standard deviation of the mean (grey vertical lines), relative to pooled data from three independent experiments, including the results of the Welch and Brown-Forsythe one-way ANOVA for significance between the indicated proteins. ****: p < 0.0001.

### ΔNp63α residues 278-302 directly bind to IMPαs but not IMPβ1

Bipartite cNLSs are characterized by the ability to bind IMPβ1 through the adapter IMPα, which has several isoforms in humans, each with specific expression patterns and NLS binding properties (74–76). We therefore tested the ability of ΔNp63α NLSbip to bind IMPβ1 as well as several IMPαΔIBB isoforms by fluorescence polarization (FP). Our results indicate that ΔNp63α NLS bound weakly to IMPβ1 (unable to determine Kd), but could be recognized by all IMPα isoforms with high affinity (K_D_’s 4.3 – 57.4 nM), with a subtle preference for IMP α1 (Kd 4.3 nM), α3 (Kd 6.7 nM) and α7 (Kd 4.6 nM) (Figure 3A). To further probe the contribution of each binding site, several ΔNp63α NLS substitution mutants were designed, replacing either the N-term (mNLSn) or C-term (mNLSc) basic amino acids with alanine. Substitution of either N-term or C-term basic amino acids strongly decreased the affinity of interaction (unable to determine K_D_), while substitution of both completely ablated binding (Figure 3B). Therefore, our data suggest that ΔNp63α is imported to the nucleus via a bipartite NLS located at residues 278-302, rather than the monopartite NLS that was described previously (71).

**Figure 3.**
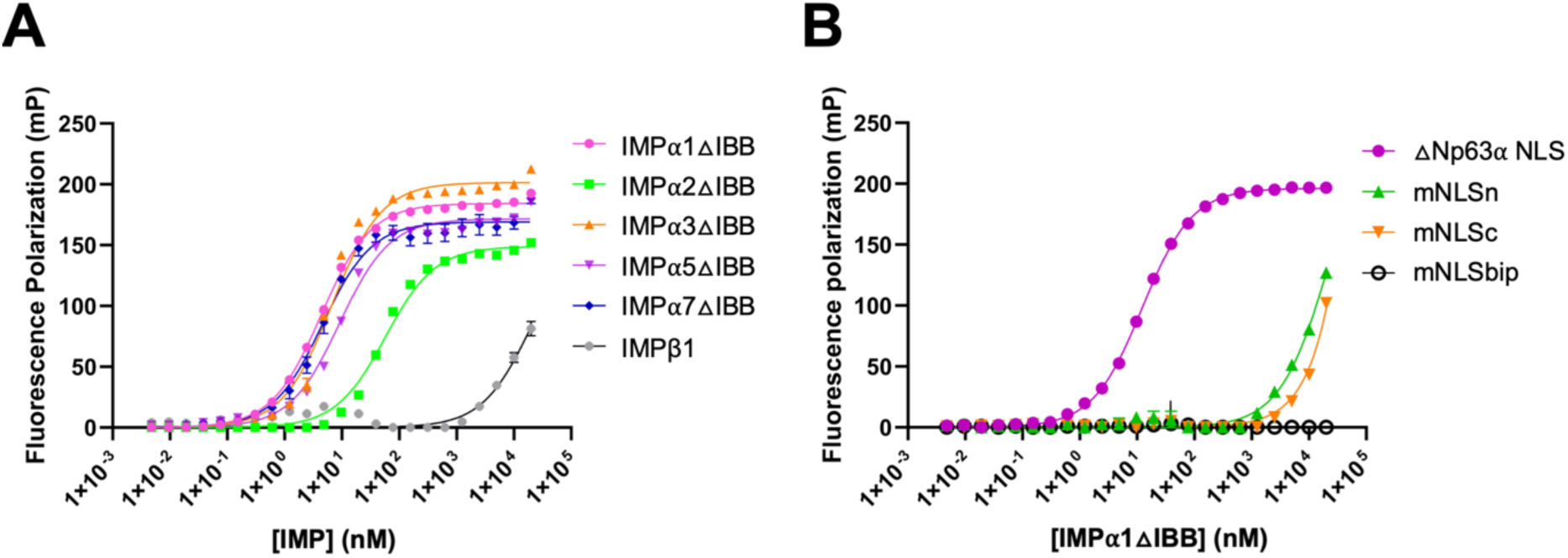
ΔNp63α residues 278-302 bind to IMPα isoforms but not to IMPβ1 with high affinity. (A) Fluorescence polarization assays measuring binding affinity between the indicated IMPs (20 µM starting concentration) and FITC-ΔNp63α NLSbip peptide (10 nM). (B) The binding of the indicated ΔNp63α NLS derivatives was tested against mIMPα1ΔIBB. Data are show as mean values ± standard error of the mean (n=3 biologically independent experiments).

### ΔNp63α residues 278-302 represent a bipartite cNLS

Since bipartite cNLSs are defined by their ability to simultaneously bind to IMPα minor and major binding sites, we utilized x-ray crystallography to resolve the structure of mIMPα2ΔIBB with ΔNp63α NLSbip (residues 278-DGTKRPFRQNTHGIQMTSIKKRRSP-302, Figure 4A). Crystals diffracted to 2.20 Å and data were indexed in the P2_1_2_1_2_1_ space group with unit cell parameters *a*=78.39, *b*=90.14 and *c*=100.08 Å (Supplementary Table S4). Analysis of the complex revealed the bipartite nature of the p63 NLS, which was bound in the IMPα2 minor binding site via NLSn (280-TKRPFR-285) and the IMP⍺2 major bind site via NLSc (297-KKRR-300) (Figure 4A). The minor site was occupied by a canonical ‘KR’ motif, provided by the NLSn Lys281 and Arg282 occupying the P1’ and P2’ sites respectively, as typically seen within other bipartite structures (42, 53, 77, 78) (Figure 4A). Accordingly, within the P1’ cavity, NLSn Lys281 forms hydrogen bonds with IMP⍺ residues Val321, Asn361, and Thr328. NLSn Arg282 occupies the P2’ cavity hydrogen bonding with IMP⍺ residues Ser360, Asn361, and Glu396 and forms a salt bridge with IMP⍺ Glu396 (Figure 4B). Within the major site of mIMPα2ΔIBB, the P1 site was occupied by ΔNp63α NLSc Ile296, hydrogen bonding with IMP⍺ Asn235. ΔNp63α NLSc Lys297 occupied the canonical P2 site forming hydrogen bonds with Thr155 and Asp192 with a salt bridge to Asp192. Within the P3 site the ΔNp63α Lys298 formed hydrogen bonds with Asn188. ΔNp63α Arg299 hydrogen bonded with IMP⍺ Arg106 and Leu104 at the P4 site. The P5 site was occupied by the ΔNp63α Arg300 forming hydrogen bonds with IMP⍺ Asn146 and Gly181 (see Supplementary Table S5 for a summary of the mIMPα2ΔIBB:ΔNp63α LSbip interactions). Therefore, we conclude ΔNp63α residues 278-DGTKRPFRQNTHGIQMTSIKKRRSP-302 represent a functional NLSbip.

**Figure 4.**
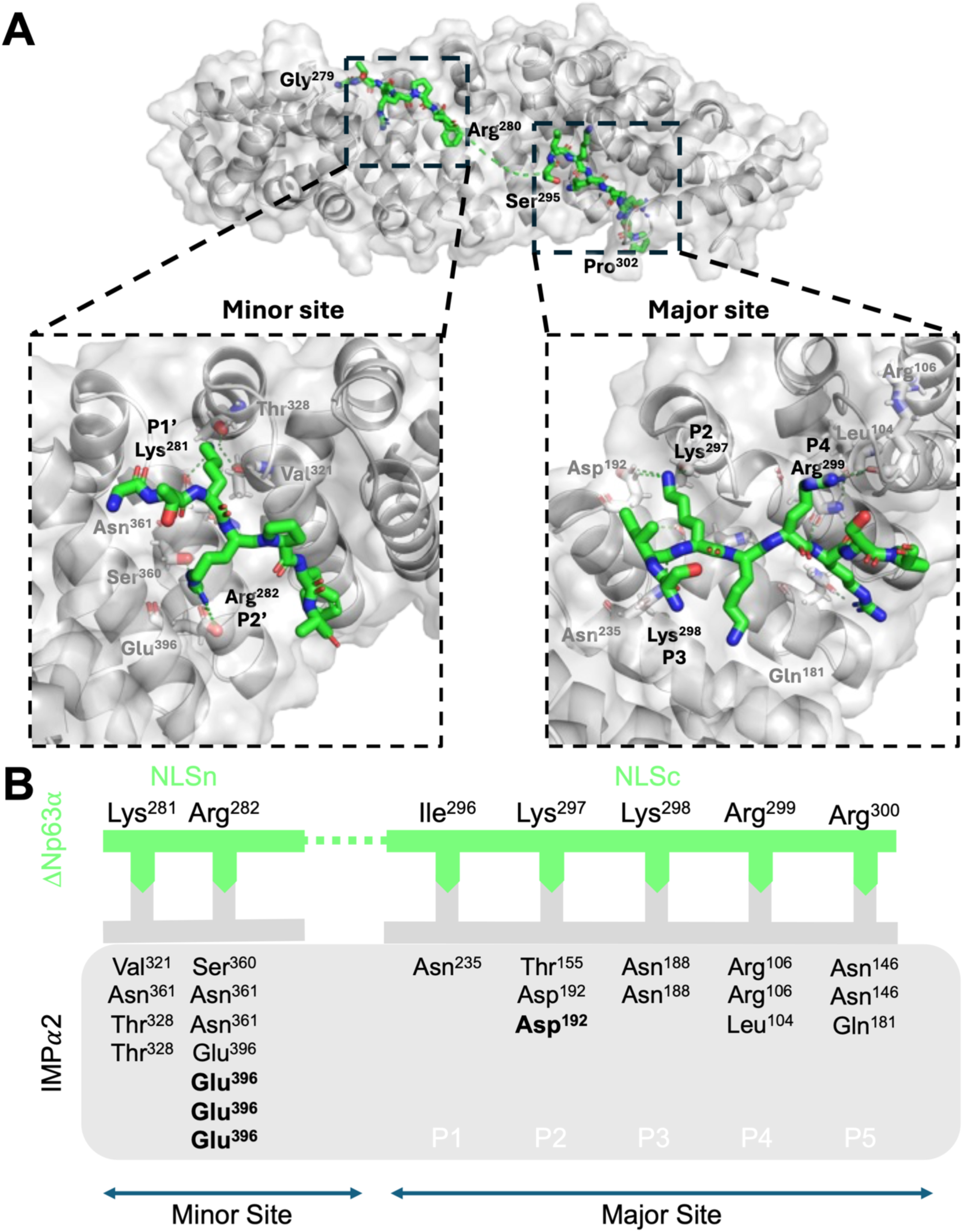
Structural characterization of ΔNp63α NLS:mIMPα2ΔIBB reveals the presence of a bipartite NLS at residues 278-302. (A) The crystal structure of FITC-labelled ΔNp63α NLS peptide was resolved in complex with mIMPα2ΔIBB to 2.20 Å resolution (PDB ID 9N54). ΔNp63α NLS is presented as green sticks bound to mIMPα2ΔIBB (grey cartoon and surface representation) bound across both the minor and major sites. ΔNp63α NLS:mIMPα2ΔIBB interactions shown in detail in boxes where mIMPα2ΔIBB interacting residues are shown as grey sticks and labels. (B) Simplified representation of ΔNp63α: mIMPα2ΔIBB binding interactions. The ΔNp63α (green line) residues interacting with mIMPα2ΔIBB (grey box) are indicated by complimentary arrows with salt bridges highlighted by bold mIMPα2ΔIBB residues as calculated using the PDBePISA server. Protein structures visualized by Pymol.

### Heterogenicity in NLS sequences across different p63 isoforms

Our biochemical, functional and structural data suggest that ΔNp63α nuclear import is mediated by a canonical NLSbip located at residues 278-302, with both basic stretches of amino acids required for high affinity interaction with IMPα and nuclear import. Importantly an analysis on the non-redundant protein database reveled that this NLSbip is highly conserved (100% identity) in the 12 best-characterized p63 human isoforms (Figure 5), and overall, in 765 different p63 isoforms from 211 mammalian species (Supplementary Table S6). However, poorly characterized p63 variants lacking the N-terminal stretch of basic amino acids have been described in human and mouse (9, 79). Such variants, often referred to as “delta”, are originated by alternative splicing of exon 9, causing the deletion of 12 base pairs and the replacement of the GTKRP pentapeptide with A, leaving the NLSbip devoid of its N-terminal stretch of basic amino acids (278-DAFRQNTHGIQMTSIKKRRSP-298, Figure 5). Despite substitution of KR to A clearly impacting both nuclear targeting and IMPα binding due to NLSbip when tested in the context of a NLS peptide (see Figure 2 and 3), the mouse isoforms characterized so far are not retained in the cytoplasm upon transient expression (9, 79). Such deletion could be identified in 311 p63 isoforms, from 164 different mammals (Figure 5 and Supplementary Table S7). Therefore, despite the NLSbip described here is highly conserved across the most-studied p63 isoforms, its N-terminal portion is deleted in the poorly characterized “delta” isoforms (also known as “DeltaExon8” (80)).

**Figure 5.**
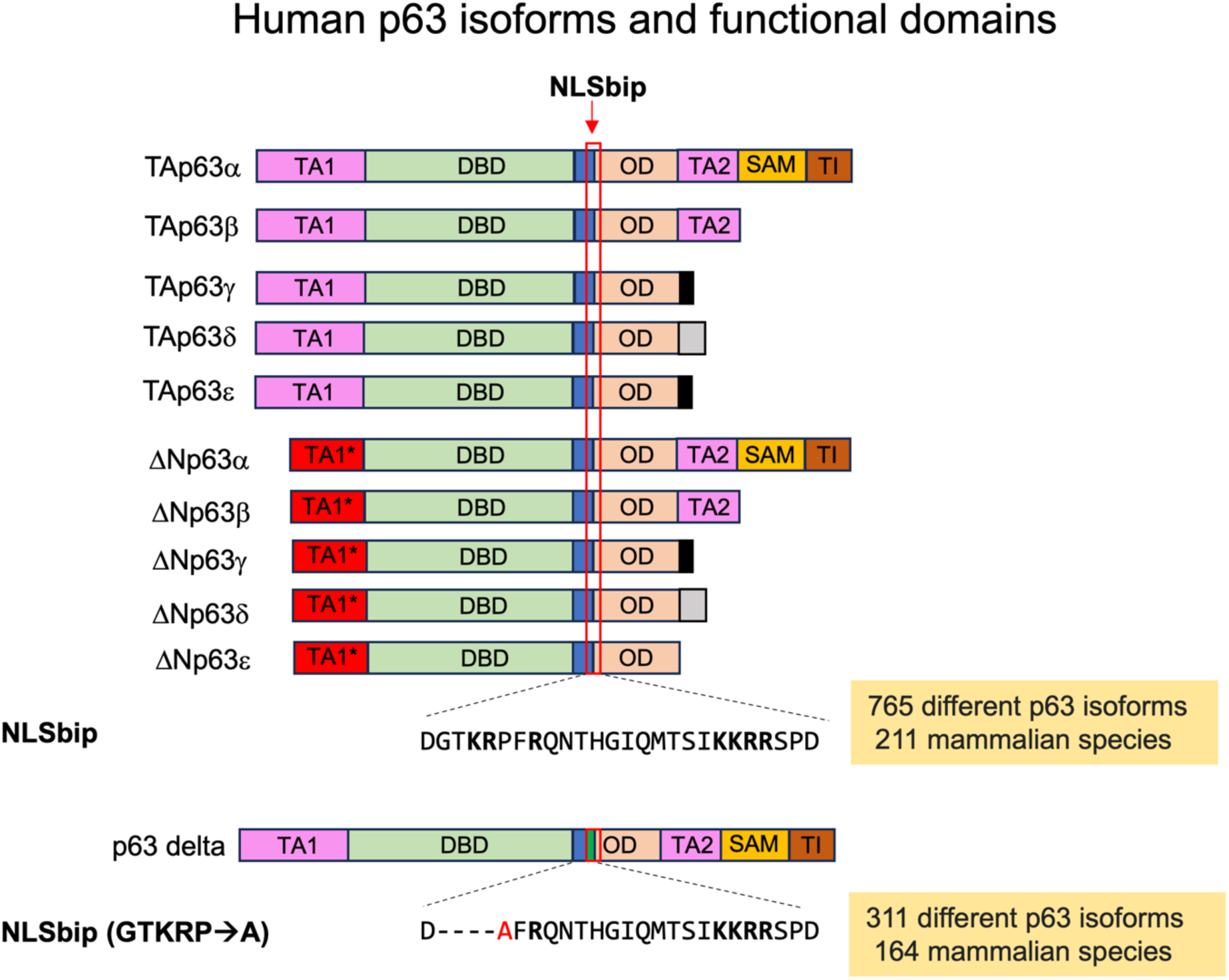
NLS heterogenicity in p63 isoforms. A schematic representation of human p63 isoforms and their functional domains. TA1: transactivator domain 1; TA1*: alternative N-terminus typical of ΔN isoforms; DBD: DNA binding domain; NLSbip: bipartite nuclear localization signal; OD: oligomerization domain; TA2: transactivation domain 2; SAM: sterile alpha motif; TI: transcriptional inhibition. The sequence of the ΔNp63α NLSbip (DGT**KR**PF**R**QNTHGIQMTSI**KKRR**SPD) was used to perform a Blast search over the non-redundant protein database. A 100% identity was observed for 765 p63 isoforms from 211 mammalian species, including the best-characterized human p63 isoforms. For 311 different p63 isoforms from 164 mammalian species, a GTKRP/A deletion within NLSn was observed, including the poorly characterized p63 delta isoform in humans.

### Nuclear import of ΔNp63α can be mediated by either the NLSn or the NLSc forming an atypical NLSbip

The evidence that p63 nuclear isoforms lacking the N-terminal stretch of basic amino acids forming NLSbip are not impaired in nuclear localization (9, 79), coupled with the report that deletion of the C-terminal basic stretch of amino acids forming NLSbip, only mildly affected nuclear localization of full length ΔNp63α (71), while their substitution to A almost completely ablated IMPα binding and nuclear targeting properties in the context of the NLSbip peptide (see Figure 2 and 3), raised the possibility that additional NLSs are present with p63 sequence, in addition to NLSbip. To verify the contribution of the latter to ΔNp63α nuclear transport, we quantified the impact of alanine substitutions of its key basic amino acids on nuclear accumulation of YFP-ΔNp63α (Figure 6A). As expected, YFP-ΔNp63α strongly accumulated in the cell nucleus (Figure 6B and Supplementary Figure S3), in 100% of analyzed cells (Figure 6C) and with an average Fn/c of 45.7 (Figure 6D). Substitution of the N-terminal NLSbip basic amino acids with alanine residues (mNLSn; K281A/R282A) affected nuclear accumulation only very mildly, resulting in a decrease of the Fn/c to 38.9 (Figure 6D). Substitution of the C-terminal NLSbip basic amino acids (mNLSc; K297A/K298A/R299A/R300A) caused a more pronounced reduction in nuclear accumulation, resulting in a protein mislocalized to the cytoplasm in c. 20% of cells (Figure 6C) and a mean Fn/c of 2.9 (Figure 6D). Intriguingly, only the substitution of both basic stretches completely ablated nuclear targeting, with YFP-ΔNp63α mNLSbip being retained in the cytosol in 99% of transfected cells (Figure 6C) and a mean Fn/c of 0.5 (Figure 6D). Therefore, each basic stretch of basic amino acids forming ΔNp63α NLSbip is sufficient to target the protein into the nucleus.

**Figure 6.**
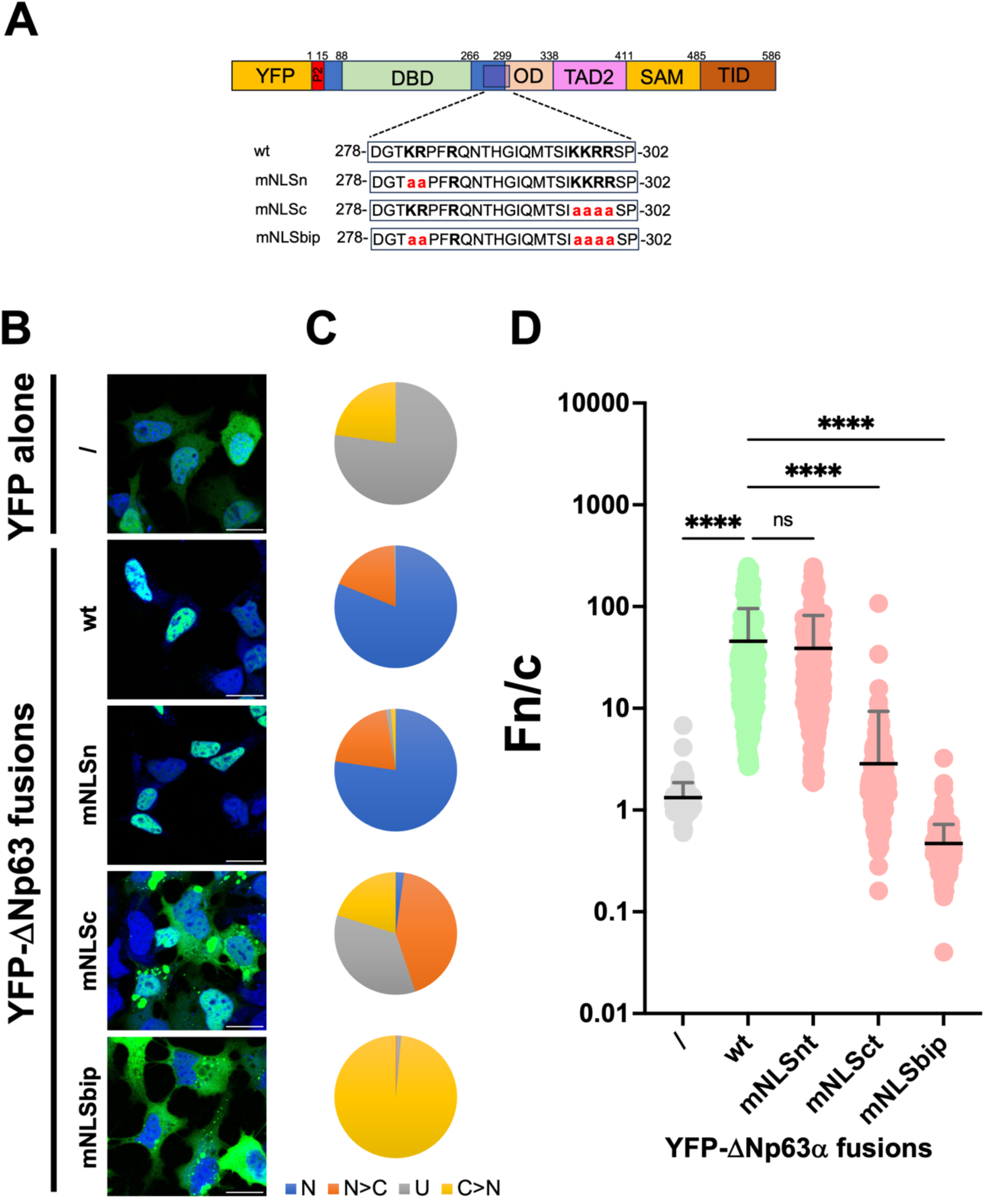
Substitution of basic residues within ΔNp63α bipartite NLS results in failure to accumulate in the cell nuclei. (A) HEK293A were transfected to express the indicated YFP-ΔNp63α fusion protein and substitution derivatives thereof. The sequence of ΔNap63 NLSbip is shown as capital letters, with basic amino acids in boldface and substituted amino acids in red lowercase. (B) Twenty-four hours later cells were incubated with DRAQ5 to stain cell nuclei, fixed, and processed for single-cell quantitative CLSM analysis as described in the Materials and Methods section. Merge images of the 633 (DRAQ5, nuclei) and 488 nm (YFP, ΔNp63α) laser lines are shown. Individual cells expressing the indicated proteins were analyzed to calculate the Fn/c value relative to YFP fusion proteins. (C) Pie charts representing the percentage of cells relative to each indicated fusion protein displaying the indicated subcellular localizations is shown. N: nuclear, Fn/c ≥ 10; N>C: nuclear more than cytosolic, 2 ≤ Fn/c < 10; U: ubiquitous, 1 ≤ Fn/c < 2; C>N: more cytosolic than nuclear, Fn/c < 1. (D) The Fn/c values relative to the indicated fusion proteins are shown as individual measurements from single cells (circles), means (black horizontal lines) and standard deviation of the mean (grey vertical lines), relative to pooled data from three independent experiments, including the results of the Welch and Brown-Forsythe one-way ANOVA for significance between the indicated proteins. ****: p < 0.0001.

### Nuclear levels of ΔNp63α regulate its transcriptional activity

Since ΔNp63α has been shown to potently inhibit p53-mediated transactivation from a minimal p53 RE by direct DNA binding (14), we reasoned that defects in nuclear import should impair prevent ΔNp63α-mediated DNA binding and transcriptional regulation. To verify this hypothesis, we developed an assay to simultaneously monitor the effect of amino acidic substitutions within ΔNp63α sequence on its DNA binding ability by measuring their impact on p53-mediated transactivation, as well as on p53-levels (Figure 7A). The assay is based on transfection of H1299 cells with plasmids mediating expression of YFP-p53 as well as RFP under control of the CMV promoter, and of NLuc under control of 2x p53 minimal RE. Such setup allows for simultaneous measurement of both transfection efficiency (RFP, see Supplementary Figure S4B), p53 amount (YFP, see Supplementary Figure S4C), and activity (NLuc, see Supplementary Figure S4D), as well as to measure p53 activity normalized either for transfection efficiency (NLuc/RFP, see Figure 7D), or for p53 expression levels (NLuc/YFP, see Figure 7E). In the absence of ΔNp63α, YFP-p53 could potently stimulate transcription (Figure 7DE). As expected, expression of FLAG-ΔNp63α strongly decreased p53 transcriptional activity in a dose-dependent fashion (Figure 7DE), without affecting YFP-p53 expression levels (Figure 7C). However, overexpression of FLAG-ΔNp63α mNLSbip had a significantly less pronounced effect on inhibition of p53-mediated transcription, indicating reduced DNA binding due to impaired nuclear localization.

**Figure 7.**
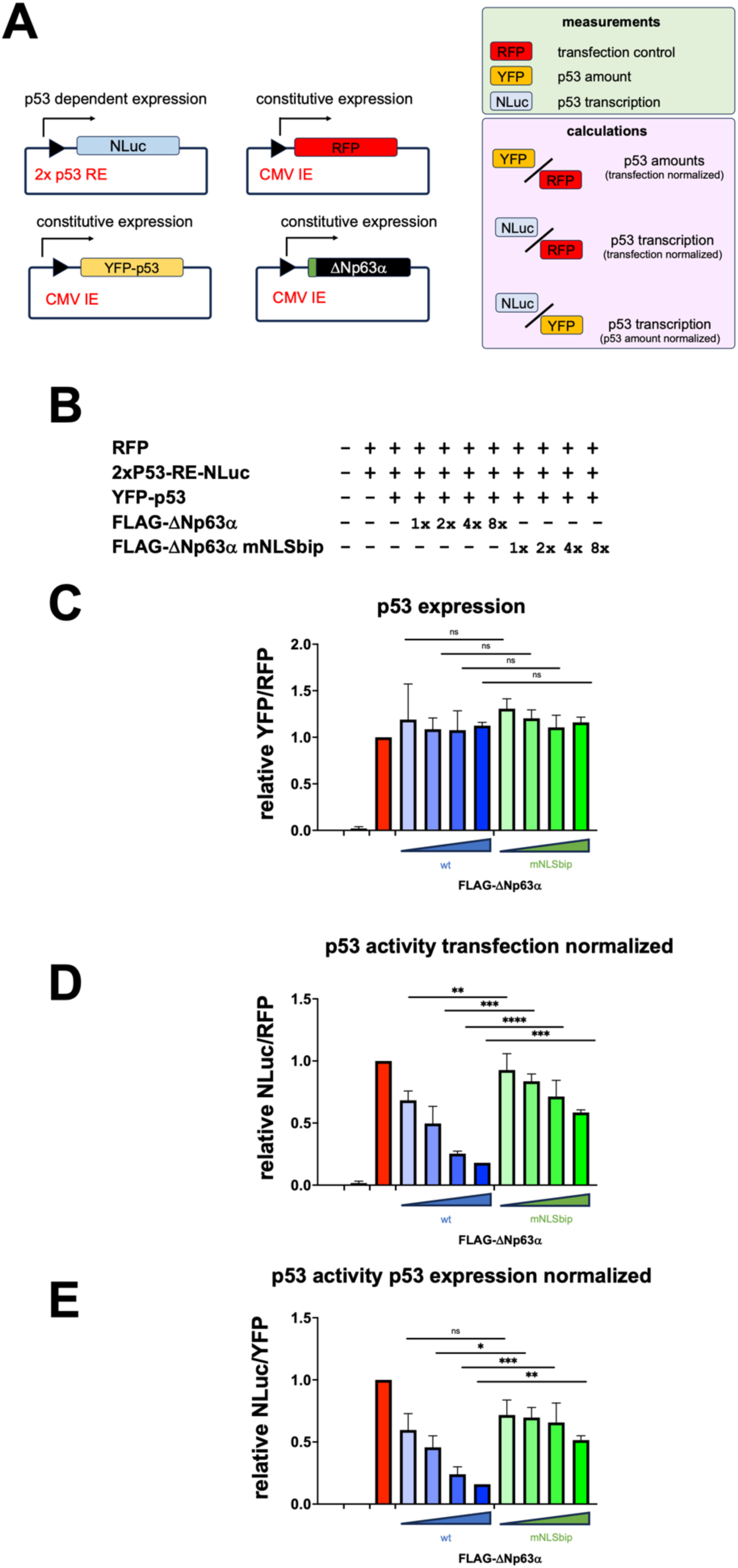
Nuclear localization is essential for ΔNp63α transcriptional inhibition. (A) Schematic representation of the p53 transcription assay. (B) H1299 were transfected with plasmids mediating expression of Tag-RFP under the control of the constitutive Immediate early (IE) promoter (CMV), to monitor transfection efficiency; nanoluciferase (NLuc) under the control of two p53 response elements (2xTP53 RE), to monitor p53-dependent transcription; YFP-p53 under the control of CMV IE promoter to transactivate 2xTP53 RE; and increasing amounts of either FLAG-ΔNp63α or FLAG-ΔNp63α;mNLSbip to repress the 2xTP53 RE. Twenty-four hours later, cells were processed as described in the Materials and Methods to perform luciferase assays and monitor 2xTP53 RE transactivation. (C) The expression of p53 under each condition and relative to cells transfected in the absence of FLAG-ΔNp63α is shown as the YFP/RFP ratio. (D) The activity of p53 under each condition, normalized per the transfection efficiency and relative to cells transfected in the absence of FLAG-ΔNp63α is shown as the NLuc/RFP ratio. (E) The activity of p53 under each condition, normalized per the levels of p53 and relative to cells transfected in the absence of FLAG-ΔNp63α is shown as the NLuc/YFP ratio. Data are shown as means + standard deviation of the mean relative to three independent experiments.

### ΔNp63α can be imported into the nucleus as a dimer

To evaluate the effect of NLS inactivating mutations on the folding of ΔNp63α, we investigated the ability of ΔNp63α NLS defective mutants to self-associate in living HEK293T cells, using endpoint BRET assays. To this end, either wild-type or NLS defective ΔNp63α derivatives were expressed as RLuc-fusions in the absence or the presence of their YFP-tagged counterparts. All YFP-(Supplementary Figure S5A) and RLuc-(Supplementary Figure S5B) ΔNp63α derivatives were expressed to similar levels. Importantly, when RLuc-ΔNp63α was expressed in the presence of YFP-ΔNp63α, luminometric emission at 535 nm was significantly increased, indicating efficient energy transfer between the two proteins (Supplementary Figure S5C). Accordingly, the BRET value measured for RLuc-ΔNp63α significantly increased in the presence of YFP-ΔNp63α (Figure 8A), resulting in a BRET ratio of 0.18±0.20 (Figure 8B). Very similar results were obtained upon expression of any ΔNp63α derivatives carrying NLS impairing substitutions, indicating their ability to efficiently self-interact (Figure 8AB). Intriguingly, the strong BRET ratio measured for ΔNp63α mNLSbip indicates that ΔNp63α can self-interact in the cytoplasm before nuclear import. To investigate this possibility, we analyzed the ability of RFP-ΔNp63α to re-localize GFP-ΔNp63α mNLSbip to the cell nucleus, when the two proteins were co-expressed in HEK293A cells. When expressed individually, RFP-ΔNp63α and GFP-Np63α mNLSbip localized to the cell nucleus and the cytoplasm, respectively (Figure 8CD). However, co-expression with RFP-ΔNp63α significantly increased the nuclear localization of GFP-ΔNp63α mNLSbip (Figure 8 CD). Therefore, the decreased transcriptional regulation properties of ΔNp63α mNLSbip depend on its reduced nuclear localization rather than defects in self-association, and ΔNp63α can self-interact in the cytoplasm before nuclear import, with ΔNp63α wt being able to piggy-back the otherwise cytoplasmic ΔNp63α mNLSbip into the nucleus.

**Figure 8.**
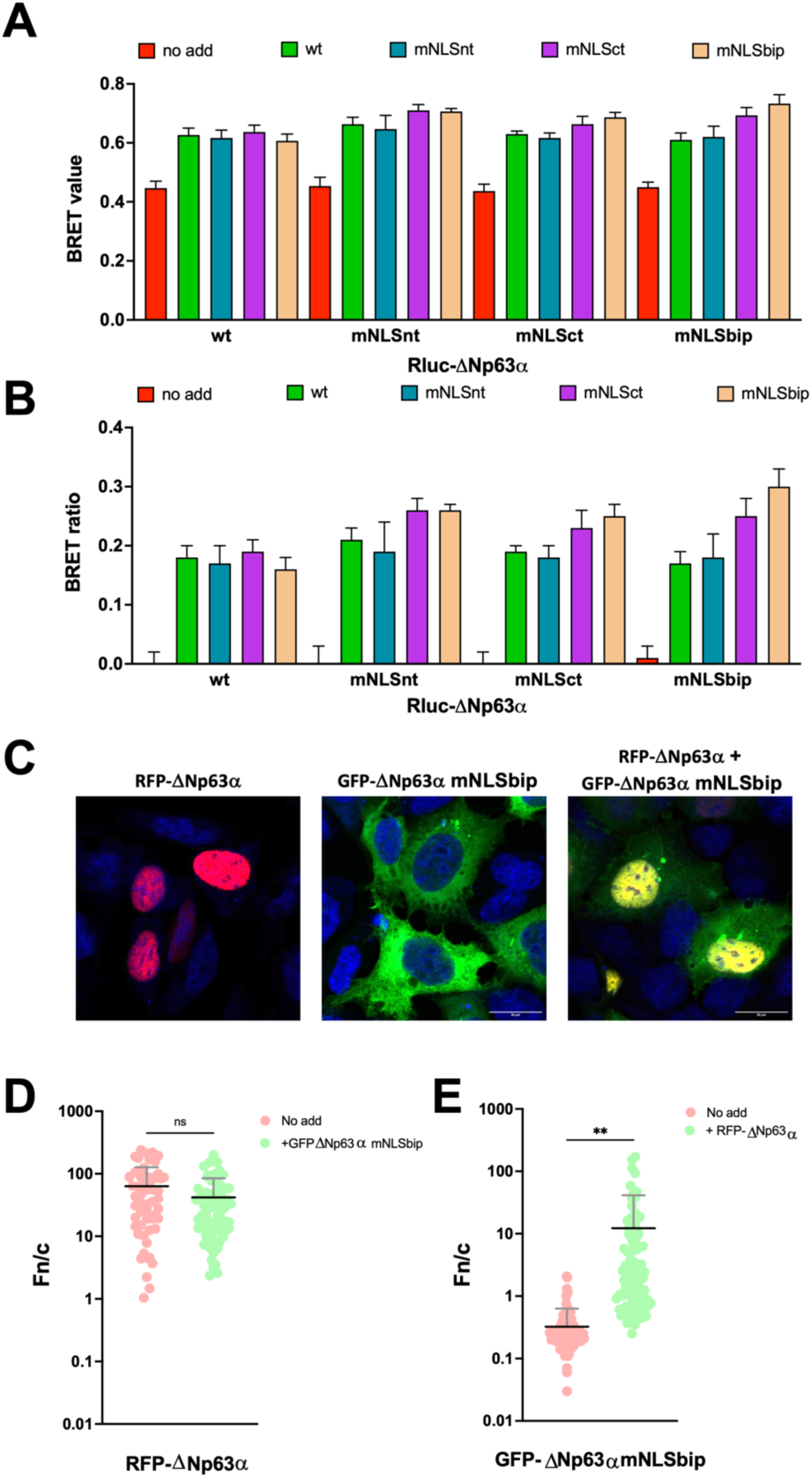
ΔNp63α can be transported in the nucleus as a dimer. (AB) HEK293T cells were transfected with 45 ng of the indicated RLuc-ΔNp63α fusion proteins in the absence (no add), or in the presence of 250 ng of plasmids mediating the expression of the indicated YFP-ΔNp63α fusion proteins. Twenty-four hours later, cells were processed as described in the Materials and Methods section to perform endpoint BRET assays, followed by calculation of the BRET value (A) and BRET ratio (B), relative to each combination. Data shown are means and standard error of the mean relative to 3 independent experiments performed in triplicate. (C-E) HEK293A were transfected to express RFP-ΔNp63α and GFP-ΔNp63α mNLSbip alone or in combination. Twenty-four hours later cells were incubated with DRAQ5 to stain cell nuclei, fixed, and processed for single-cell quantitative CLSM analysis as described in the Materials and Methods section. (C) Merge images of the 633 (DRAQ5, nuclei), 561 (RFP, ΔNp63α), and 488 nm (GFP, ΔNp63α mNLSbip) laser lines are shown. Individual cells expressing the indicated proteins were analyzed to calculate the Fn/c value relative to fusion proteins. (D) The Fn/c values relative to RFP-ΔNp63α when expressed alone (pink) or in combination with GFP-ΔNp63α mNLSbip (green) are shown as individual measurements from single cells (circles), means (black horizontal lines) and standard deviation of the mean (grey vertical lines), relative to pooled data from two independent experiments. (E) The Fn/c values relative to GFP-ΔNp63α mNLSbip when expressed alone (pink) or in combination with RFP-ΔNp63α mNLSbip (green) are shown as individual measurements from single cells (circles), means (black horizontal lines) and standard deviation of the mean (grey vertical lines), relative to pooled data from two independent experiments, including the results of the Welch and Brown-Forsythe one-way ANOVA for significance between the indicated proteins. **: p < 0.01.

### A mutation resistant three-dimensional bipartite NLS allows nuclear transport of ΔNp63α homodimers

The finding that ΔNp63α can self-interact in the cytoplasm prior to nuclear import offers a potential explanation for the apparent discrepancy between results obtained using the isolated ΔNp63α NLS and the full-length protein. Specifically, while individual mutations in either the N-terminal (NLSn) or C-terminal (NLSc) NLS motifs have minimal impact on the nuclear localization of full-length ΔNp63α (Figure 6), they strongly impair both nuclear import (Figure 2) and IMPα binding (Figure 3) when tested in the context of an isolated ΔNp63α bipartite NLS peptide. The latter observation is consistent with the known requirement of bipartite NLSs to simultaneously engage both the major and minor binding sites of IMPα for efficient nuclear transport (29). We hypothesized that ΔNp63α dimers might enable such bipartite interactions in trans, similar to the recently described p50/p65 complex, in which the bipartite NLS spans two subunits (81). To explore this, we used AlphaFold to model ΔNp63α_278-342_ dimers, encompassing the NLS through to the tetramerization domain, bound to mIMPα2ΔIBB. In the wild-type dimer (wt+wt), the bipartite NLS was presented as a conventional, intramolecular motif derived from a single monomer (Figure 9A). Interestingly, when both monomers carried mutations in the N-terminal NLS (mNLSn+mNLSn), the C-terminal NLSs occupied both the major and minor binding sites on IMPα (Figure 9B). Conversely, when both monomers had mutations in the C-terminal NLS (mNLSc+mNLSc), the N-terminal NLSs occupied both binding sites (Figure 9C). We also examined a mixed dimer with N- and C-NLS mutations in one monomer and wild-type NLSs in the other (mNLSbip+wt). This configuration resembled the wild-type complex. Together, these findings suggest that in the native context, ΔNp63α likely uses a canonical bipartite NLS configuration. However, for isoforms lacking NLSn, or in the presence of NLS mutations, dimerization may enable compensatory trans-binding, allowing the NLS elements from each monomer to cooperate in engaging IMPα and facilitating nuclear import.

**Figure 9.**
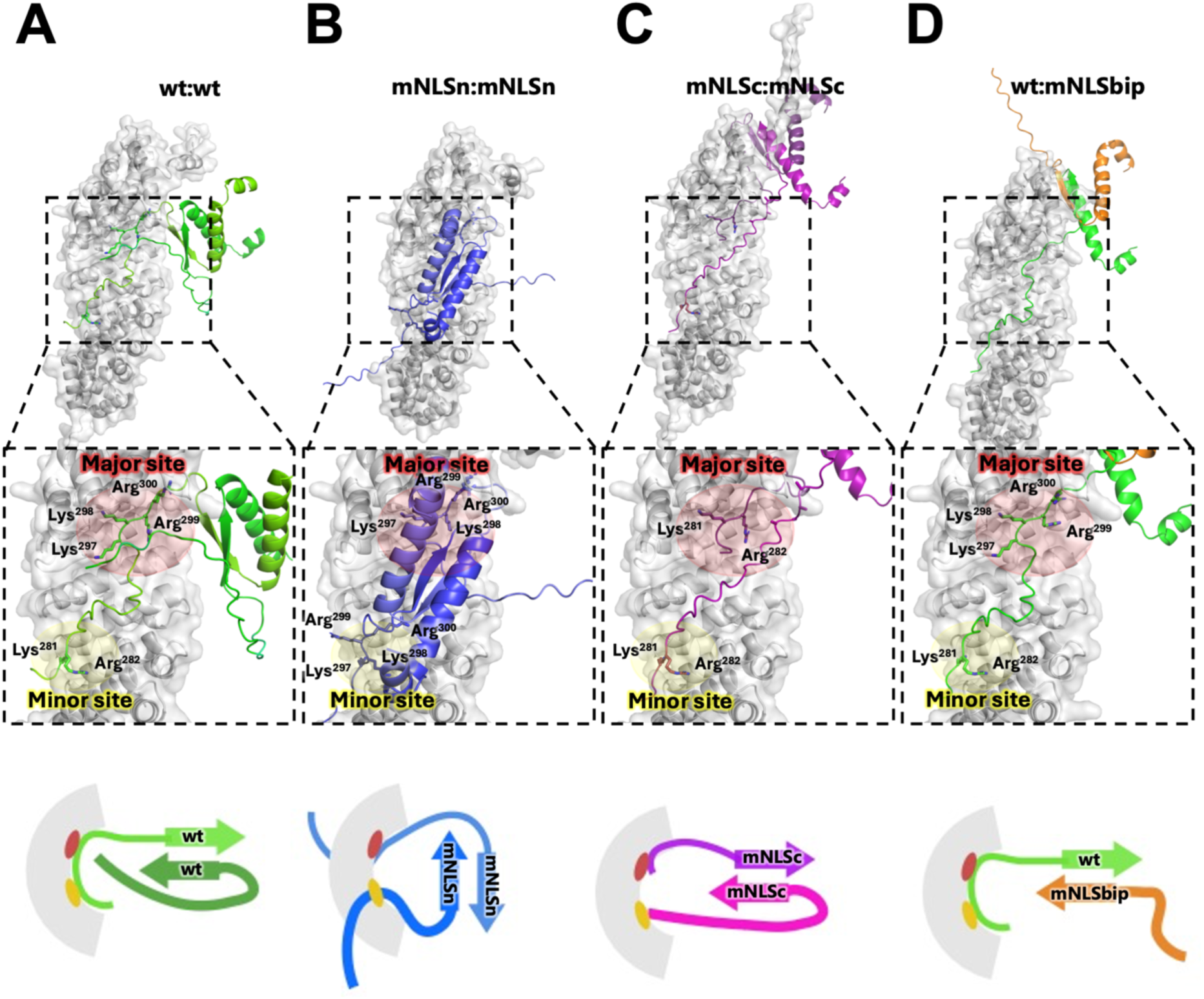
ΔNp63α forms homodimers which can rescue the mNLSn and mNLSc nuclear import in trans. Structural models were generated using AlphaFold 3 (for dimers of ΔNp63α_278-342_ (278- DGT**KR**PFRQNTHGIQMTSI**KKRR**SPDDELLYLPVRGRETYEMLLKIKESLELMQYLPQHTIETYR-342), encompassing the NLS and oligomerization domain, in complex with mIMP⍺2. Top panels show full models with close-up views of the NLS residues engaging the major (red circle) and minor (yellow circle) sites of mIMP⍺2. (A) Wt dimer of ΔNp63α_278-342,_ green cartoon with NLSn and NLSc residues shown as sticks, binds mIMP⍺2 in a bipartite manner with a single molecule occupying both minor and major sites. (B) ΔNp63α_278-342_ mNLSn dimer, purple cartoon with NLSc shown as sticks, binds the mIMP⍺2 with a C-terminal NLS from each monomer. (C) ΔNp63α_278-342_ mNLSc dimer, purple cartoon with NLSn shown as sticks, binds the mIMP⍺2 with a N-terminal NLS from each monomer. (D) The ΔNp63α_278-342_: ΔNp63α_278-342_ mNLSbip heterodimer bound the mIMP⍺2 in a bipartite manner with the wt monomer occupying both major and minor sites.

### Substitutions within ΔNp63α NLS found in the human population are not expected to impair nuclear localization

Given the importance of ΔNp63α nuclear targeting for its transcriptional activity, the tolerance of its NLSbip to mutations likely offers several evolutionary advantages over the standard scenario whereby both basic stretches of amino acids forming a NLSbip are required for cargo nuclear localization. Indeed, mutations that simultaneously target both p63 NLSbip basic stretches and prevent nuclear import should be very rare and potentially associated with pathogenic conditions. Accordingly, such p63 variants are not described in the literature. However, a few mutations affecting individual residues within NLSbip are reported in the GnomAD v4.1.0 database (69), although with very low frequency. To investigate this further, we performed a bioinformatic analysis using the VarSome and Franklin by genoox platforms to evaluate their potential clinical significance based on in silico prediction algorithms (69, 70). Furthermore, the potential impact on NLS activity was predicted by cNLS mapper (38) and based on our experimental data. Intriguingly, 30 missense mutations, affecting 17 different residues were reported (Figure 10). Among the latter, four mutations affect residues involved in interaction with IMPα (K281, R282 and R285 in the minor site, and R299 in the major site). Importantly the MAF in the European cohort for such mutations was very low (range 1: 1:32’702-1: 111’920). Furthermore, despite the K281N, R282C and R282H substitutions were predicted by cNLS mapper to disrupt the activity of the NLSbip identified here, our finding that ΔNp63α NLSc is sufficient to mediate nuclear import of the full-length protein (Figure 6), is consistent with ΔNp63α NLS activity being preserved. Additionally, the single mutation of a basic residue interacting with IMPα major site involves R299, which is accommodated in the P4 position (Figure 4), with a low thermodynamic contribution, and it is therefore extremely unlikely to cause a drastic reduction of nuclear import, unless combined with further mutations affecting other NLS residues (82). Most of the identified mutations were predicted to be of an uncertain clinical significance (US), seven yielded a controversial result, intermediate between US and likely pathogenetic (LP), two were predicted to be likely benign (LB), and only R285C was predicted be LP (Figure 10). Therefore, despite a few mutations have been described within ΔNp63α NLSbip identified here, none is expected to preclude its nuclear import.

**Figure 10.**
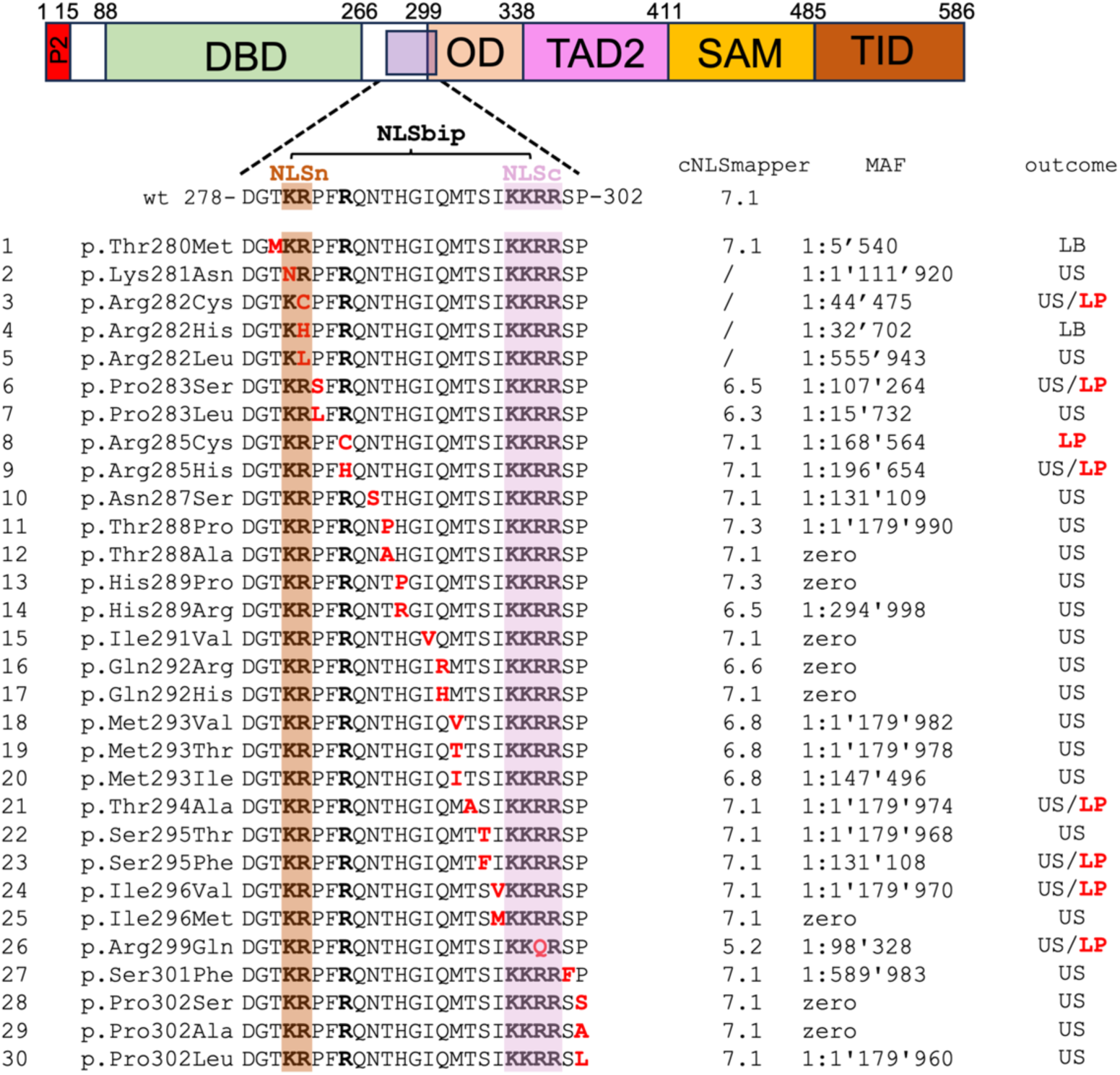
Mutations described in the human population found in ΔNp63α NLS domain are not expected to impair nuclear localization. TP63 variants were obtained from the GnomAD v4.1.0 database. Residue numbers are referred to the ΔNp63α isoform. The Minor allele frequency (MAF) in the European cohort is shown, along with the pathogenic potential according to VarSome and Franklin, the cNLS mapper score, and the predicted subcellular localization based on our experimental data. US: uncertain significance; LP: likely pathogenic; LB: Likely Benign; N: nuclear. ΔNp63α functional domains are schematically represented along with the NLSbip sequence using the single-letter amino acid code. Basic residues accommodated in the minor and major IMPα2 sites are highlighted. Basic residues are in boldface; mutations are in red.

## DISCUSSION

We have extensively characterized the nuclear import pathway of ΔNp63α, identifying its NLS and the cellular transporters involved. The process is entirely dependent on the IMPα/β1 heterodimer (Figure 1) and on two basic stretches of amino acids located at residues 278-302 (Figure 2B), forming a NLSbip apparently reminiscent of that originally described on Nucleoplasmin (Figure 4; (83)), but which is functionally tolerant to mutations targeting either the upstream or downstream stretch of basic amino acids (Figure 6). BRET and CLSM data, showing that ΔNp63α can self-interact in the cytoplasm before nuclear import (Figure 8), are consistent with our structural prediction models, showing that when NLS elements are mutated, ΔNp63α dimerization can compensate for impaired NLS motifs by forming trans-acting NLS configurations (Figure 9). This would enable robust nuclear transport even in the presence of single-site disruptions typical of specific ΔNp63α isoforms (Figure 5) or caused by rare mutations (Figure 10). The bipartite nature of ΔNp63α NLS is supported by a wide range of functional (Figure 2), biochemical (Figure 3), and structural (Figure 4) assays. Overall, ΔNp63α NLSbip comprises a newly identified stretch of basic amino acids (NLSn: 278-DGTKRPFRQ-285) located upstream of a NLS previously described (NLSc: SIKKRRSPDD-304), but whose inactivation was not sufficient to prevent nuclear import (71). Therefore, this is the first report of a completely mislocalized p63 mutant derivative and allowed an unambiguous study of the relationship between protein function and localization (see Figures 6-7). Furthermore, this is the first structural analysis to date of the complex between IMPa, and the NLS from a p53 family member (Figure 4). Our structural analysis identified DGTKRPFRQ-285 at IMPa2 minor binding site and residues SIKKRRSPDD-304 at the major site, closely resembling the interaction of other well-characterized bipartite NLSs with IMPα (53). Despite lacking structural data for p53 and p73, biochemical and cellular data are consistent with the presence of a bipartite NLS in all p53 family members (31, 36). Our study represents also the first formal proof that p53 family members are imported through the IMPa/β1 dependent pathway. Indeed, the well-characterized IMPa/β1 inhibitor Bimax2 completely ablated its nuclear import (Figure 1). Several studies have shown that Bimax2 can selectively impair the nuclear import of cargoes depending on direct interaction with IMPa, such as HCMV-UL44 (67), Parvovirus B19 nonstructural protein 1 (44), Large Tumor Antigens from the Polyomavirus family (53), but not of proteins relying on alternative IMPs such as IMPβ1 (84) or transportin (85). Overall, ΔNp63α NLS appears highly similar to that previously identified on p53 (31) and p73 (36), but presents peculiar distinguishing properties. First of all, ΔNp63α NLSbip appears endowed with a broader IMPα isoform binding ability than that of p53. Indeed, p53 has been shown to specifically interact with IMPa3, but not other IMPs (34, 86), whereas ΔNp63α NLSbip could bind both IMPα1, 3, and 7 with very similar affinity (Figure 3). This might relate to its constitutive nuclear localization, in contrast with the ability of p53 to mainly localize in the cytosol in the absence of stimuli, dependent on the ubiquitination of its NLS (87). Secondly, within its physiological context the NLSbip of ΔNp63α appears more tolerant to substitutions of its N- and C-terminal basic stretch of amino acids as compared to that of p53 and other NLSbip, whereby mutations of single residues are sufficient to strongly impair nuclear accumulation (see Table I and references therein). Indeed, simultaneous inactivation of both ΔNp63α basic stretches of amino acids is required to prevent nuclear import (Figure 6). This is the consequence of the ability of ΔNp63α to self-interact in the cytoplasm before nuclear import (Figure 8) and of the unusual three-dimensional flexibility of the ΔNp63α bipartite NLS, which allows each basic stretch to interact with both IMPα binding sites (Figure 9). This structural feature likely provides two key evolutionary advantages. First, it enables nuclear localization of ΔNp63α isoforms with divergent NLS sequences generated by alternative splicing events (9, 79, 80)— including human p63 delta (Q9H3D4-11), mouse TA*p63γ (O88898-5) and ΔNp63γ (O88898-6), as well as multiple zebrafish isoforms (A0AB32TT28, A0A8M1PDU3, Q8JFE3)— which bear the GTKRP to A change within NLSn (Figure 5 and Supplementary Table S7). Second, it increases tolerance to de novo mutations that might otherwise impair nuclear import. Consistently, GnomAD v4.1.0 reports only rare mutations in TP63 NLSbip residues, affecting individual basic amino acids—K281 and R282 (minor site), and R299 (major site)—which, based on our structural and functional data, are not expected to compromise import efficiency (Figures 4, 8, and 10). These findings suggest that mutation-tolerant bipartite NLSs, like that of ΔNp63α, may have evolved in critical proteins to reduce the impact of sequence variation on nuclear localization, particularly during development. Finally, we investigated the correlation between ΔNp63a nuclear localization, tetramerization, and transcriptional activity. Our data shows NLS defective, cytoplasmic restricted ΔNp63α can functionally self-interact (Figure 8), however, nuclear localization is required for ΔNp63αDNA binding and transcriptional repression of p53 (Figure 7). This finding is consistent with the evidence that knockdown of Nup62, which is involved in ΔNp63α nuclear transport, altered transcription of several ΔNp63α target genes in SCC cells (71), and the well-established correlation between p53 and p73 localization and activity, with decreased nuclear localization correlating with decreased transcriptional activity (18, 31, 34, 35, 86, 88).

**Table I.**
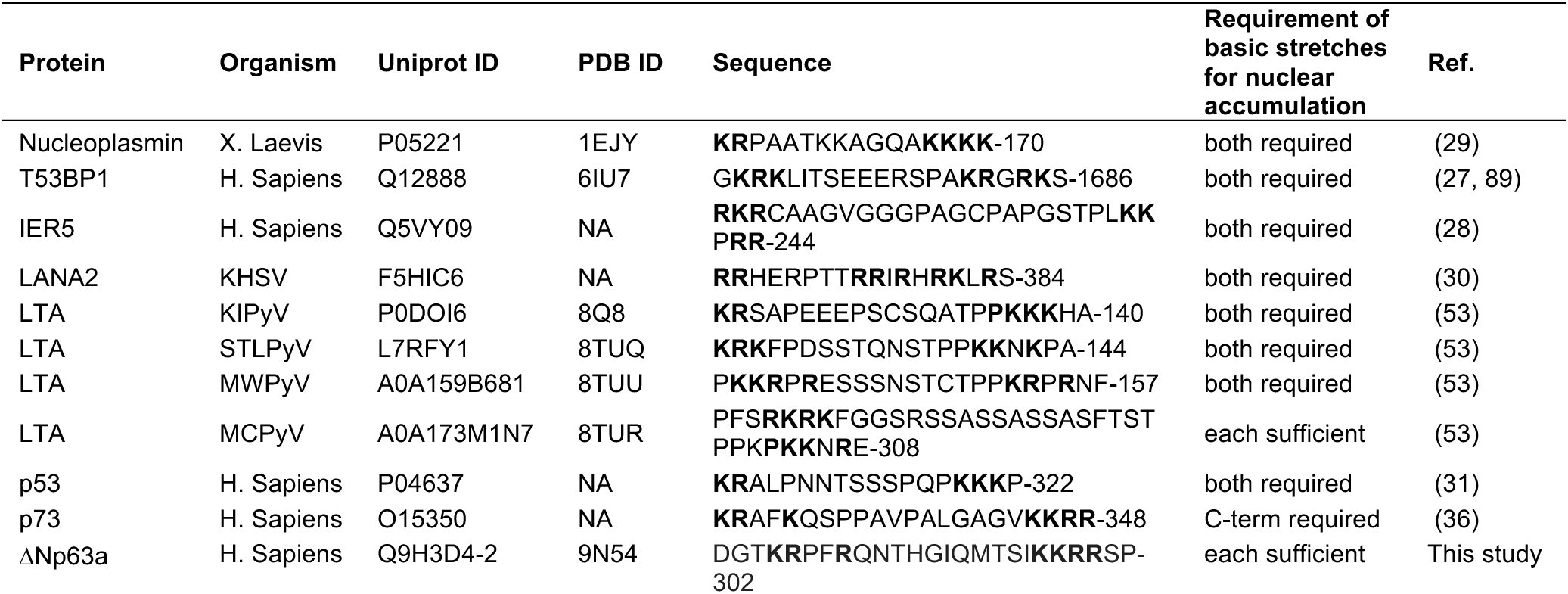
Most bipartite NLSs require both basic stretches of amino acids to mediate nuclear import. A list of well-characterized bipartite NLSs, along with their sequence, the Uniprot ID of the protein carrying them, and the PDB ID of complexes with IMPα. NA: not available.

## Supporting information

Supplementary Figurs S1-S5 and SUpplementary Tables S1-S6

## DATA AVAILABILITY

The original data presented in the study are openly available in Research data UNIPD at https://researchdata.cab.unipd.it/ (to be filled in during proofs).

## SUPPLEMENTARY DATA

The following Supplementary data is available online.: Figure S1. ΔNp63α is transported into the nucleus by the IMPα/β1 heterodimer. Supplementary Figure S2. ΔNp63α residues 278-302 represent a functional bipartite NLS. Supplementary Figure S3. Substitution of basic residues within ΔNp63α bipartite NLS results in failure to accumulate in the cell nuclei. Supplementary Figure S4. Nuclear localization is essential for ΔNp63α transcriptional inhibition. Raw data relative to Figure 7. Supplementary Figure S5. Nuclear localization is not required for ΔNp63α self-association. Supplementary Table S1. Plasmids used in this study. Supplementary Table S2. Sequences of peptides used in this study. Supplementary Table S3. Summary Statistics of FP assays shown in Figure 3. Supplementary Table S4. Data collection and refinement statistics for structure of ΔNp63α NLS in complex with IMPα. Supplementary Table S5. Hydrogen bonds and salt bridges of ΔNp63α NLS:mIMPα2ΔIBB complex. Supplementary Table S6. BlastP hits with 100% identify to human ΔNp63α NLSbip (DGTKRPFRQNTHGIQMTSIKKRRSP). Supplementary Table S7. BlastP hits with 100% identify to human ΔNp63α NLS (DAFRQNTHGIQMTSIKKRRSP).

## AUTHOR CONTRIBUTIONS

Gualtiero Alvisi: Conceptualization, Formal analysis, Data Curation, Methodology, Validation, Visualization, Supervision, Writing—original draft, Funding acquisition. Anna De Marinis: Investigation, Visualization, Formal analysis. Sara Esmaeili: Investigation, Formal analysis, Validation, Visualization, Writing – review and editing. Simone Ranno: Investigation, Visualization. Silvia Pavan: Investigation, Visualization. Crystall Swarbrick: Investigation, Formal analysis, Supervision, Validation, Visualization, Writing – review and editing. Brian McSharry: Formal analysis, Validation, Supervision, Writing – review and editing. Jade Forwood: Conceptualization, Formal analysis, Data Curation, Methodology, Validation, Visualization, Supervision, Writing – review and editing, Funding acquisition. Martin Pal: Formal analysis, Validation, Supervision, Writing – review and editing. Enzo Di Iorio: Funding acquisition, Resources, Writing – review and editing.

## ACKNOWLEDGEMENTS

We thank Arianna Loregian (Padova, Italy) for providing H1299 cells, Yoshihiro Yoneda and Masahiro Oka (Osaka, Japan) for gifting plasmid mCherry-Bimax2, Roger Tsien (San Diego, USA) for plasmid Plasmid Tag-RFP-N, Pietro Scaturro (Hamburg) for pDESTntYFP, pDESTntRLuc, Koen Venken (Ghent, Belgium) for plasmid 2xP53_RE::NLuc, and Emanuele Panza (Bologna) for plasmid pDESTntFLAG. This research was undertaken in part using the MX2 beamline at the Australian Synchrotron, part of ANSTO, and made use of the Australian Cancer Research Foundation (ACRF) detector.

## REFERENCES

1. Barbieri CE, Pietenpol JA. p53 family members: similar biochemistry, different biology. Cancer biology & therapy. 2005;4(4):419–20.

2. Stiewe T. The p53 family in differentiation and tumorigenesis. Nat Rev Cancer. 2007;7(3):165–8.

3. Barbieri CE, Pietenpol JA. p63 and epithelial biology. Experimental cell research. 2006;312(6):695–706.

4. Yang A, Schweitzer R, Sun D, Kaghad M, Walker N, Bronson RT, et al. p63 is essential for regenerative proliferation in limb, craniofacial and epithelial development. Nature. 1999;398(6729):714-8.

5. Petitjean A, Ruptier C, Tribollet V, Hautefeuille A, Chardon F, Cavard C, et al. Properties of the six isoforms of p63: p53-like regulation in response to genotoxic stress and cross talk with DeltaNp73. Carcinogenesis. 2008;29(2):273–81.

6. Arrowsmith CH. Structure and function in the p53 family. Cell Death Differ. 1999;6(12):1169–73.

7. Murray-Zmijewski F, Lane DP, Bourdon JC. p53/p63/p73 isoforms: an orchestra of isoforms to harmonise cell differentiation and response to stress. Cell Death Differ. 2006;13(6):962–72.

8. Mangiulli M, Valletti A, Caratozzolo MF, Tullo A, Sbisa E, Pesole G, et al. Identification and functional characterization of two new transcriptional variants of the human p63 gene. Nucleic Acids Res. 2009;37(18):6092–104.

9. Yang A, Kaghad M, Wang Y, Gillett E, Fleming MD, Dotsch V, et al. p63, a p53 homolog at 3q27-29, encodes multiple products with transactivating, death-inducing, and dominant-negative activities. Mol Cell. 1998;2(3):305–16.

10. Mills AA, Zheng B, Wang XJ, Vogel H, Roop DR, Bradley A. p63 is a p53 homologue required for limb and epidermal morphogenesis. Nature. 1999;398(6729):708-13.

11. Keyes WM, Pecoraro M, Aranda V, Vernersson-Lindahl E, Li W, Vogel H, et al. DeltaNp63alpha is an oncogene that targets chromatin remodeler Lsh to drive skin stem cell proliferation and tumorigenesis. Cell Stem Cell. 2011;8(2):164–76.

12. Yang A, Zhu Z, Kapranov P, McKeon F, Church GM, Gingeras TR, et al. Relationships between p63 binding, DNA sequence, transcription activity, and biological function in human cells. Mol Cell. 2006;24(4):593–602.

13. Candi E, Rufini A, Terrinoni A, Dinsdale D, Ranalli M, Paradisi A, et al. Differential roles of p63 isoforms in epidermal development: selective genetic complementation in p63 null mice. Cell Death Differ. 2006;13(6):1037–47.

14. Bougeard G, Hadj-Rabia S, Faivre L, Sarafan-Vasseur N, Frebourg T. The Rapp-Hodgkin syndrome results from mutations of the TP63 gene. Eur J Hum Genet. 2003;11(9):700–4.

15. Natan E, Joerger AC. Structure and kinetic stability of the p63 tetramerization domain. J Mol Biol. 2012;415(3):503–13.

16. Russo C, Osterburg C, Sirico A, Antonini D, Ambrosio R, Wurz JM, et al. Protein aggregation of the p63 transcription factor underlies severe skin fragility in AEC syndrome. Proc Natl Acad Sci U S A. 2018;115(5):E906–E15.

17. Novelli F, Ganini C, Melino G, Nucci C, Han Y, Shi Y, et al. p63 in corneal and epidermal differentiation. Biochem Biophys Res Commun. 2022;610:15–22.

18. O’Brate A, Giannakakou P. The importance of p53 location: nuclear or cytoplasmic zip code? Drug resistance updates : reviews and commentaries in antimicrobial and anticancer chemotherapy. 2003;6(6):313–22.

19. Hakhverdyan Z, Molloy KR, Keegan S, Herricks T, Lepore DM, Munson M, et al. Dissecting the Structural Dynamics of the Nuclear Pore Complex. Mol Cell. 2021;81(1):153–65 e7.

20. Timney BL, Raveh B, Mironska R, Trivedi JM, Kim SJ, Russel D, et al. Simple rules for passive diffusion through the nuclear pore complex. The Journal of cell biology. 2016;215(1):57–76.

21. Wing CE, Fung HYJ, Chook YM. Karyopherin-mediated nucleocytoplasmic transport. Nat Rev Mol Cell Biol. 2022;23(5):307–28.

22. Kimura M, Morinaka Y, Imai K, Kose S, Horton P, Imamoto N. Extensive cargo identification reveals distinct biological roles of the 12 importin pathways. eLife. 2017;6.

23. Mackmull MT, Klaus B, Heinze I, Chokkalingam M, Beyer A, Russell RB, et al. Landscape of nuclear transport receptor cargo specificity. Molecular systems biology. 2017;13(12):962.

24. Lange A, Mills RE, Lange CJ, Stewart M, Devine SE, Corbett AH. Classical nuclear localization signals: definition, function, and interaction with importin alpha. J Biol Chem. 2007;282(8):5101–5.

25. Conti E, Uy M, Leighton L, Blobel G, Kuriyan J. Crystallographic analysis of the recognition of a nuclear localization signal by the nuclear import factor karyopherin alpha. Cell. 1998;94(2):193–204.

26. Lange A, McLane LM, Mills RE, Devine SE, Corbett AH. Expanding the definition of the classical bipartite nuclear localization signal. Traffic. 2010;11(3):311–23.

27. von Morgen P, Lidak T, Horejsi Z, Macurek L. Nuclear localisation of 53BP1 is regulated by phosphorylation of the nuclear localisation signal. Biol Cell. 2018;110(6):137–46.

28. Yamano S, Kimura M, Chen Y, Imamoto N, Ohki R. Nuclear import of IER5 is mediated by a classical bipartite nuclear localization signal and is required for HSF1 full activation. Experimental cell research. 2020;386(1):111686.

29. Robbins J, Dilworth SM, Laskey RA, Dingwall C. Two interdependent basic domains in nucleoplasmin nuclear targeting sequence: identification of a class of bipartite nuclear targeting sequence. Cell. 1991;64(3):615–23.

30. Munoz-Fontela C, Rodriguez E, Nombela C, Arroyo J, Rivas C. Characterization of the bipartite nuclear localization signal of protein LANA2 from Kaposi’s sarcoma-associated herpesvirus. The Biochemical journal. 2003;374(Pt 2):545–50.

31. Liang SH, Clarke MF. A bipartite nuclear localization signal is required for p53 nuclear import regulated by a carboxyl-terminal domain. J Biol Chem. 1999;274(46):32699–703.

32. Shaulsky G, Goldfinger N, Ben-Ze’ev A, Rotter V. Nuclear accumulation of p53 protein is mediated by several nuclear localization signals and plays a role in tumorigenesis. Mol Cell Biol. 1990;10(12):6565–77.

33. Ikliptikawati DK, Makiyama K, Hazawa M, Wong RW. Unlocking the Gateway: The Spatio-Temporal Dynamics of the p53 Family Driven by the Nuclear Pores and Its Implication for the Therapeutic Approach in Cancer. International journal of molecular sciences. 2024;25(13).

34. Marchenko ND, Hanel W, Li D, Becker K, Reich N, Moll UM. Stress-mediated nuclear stabilization of p53 is regulated by ubiquitination and importin-alpha3 binding. Cell Death Differ. 2010;17(2):255–67.

35. Hara T, Arai K, Koike K. Form of human p53 protein during nuclear transport in Xenopus laevis embryos. Experimental cell research. 2000;258(1):152–61.

36. Inoue T, Stuart J, Leno R, Maki CG. Nuclear import and export signals in control of the p53-related protein p73. J Biol Chem. 2002;277(17):15053–60.

37. Borlido J, D’Angelo MA. Nup62-mediated nuclear import of p63 in squamous cell carcinoma. EMBO reports. 2018;19(1):3–4.

38. Kosugi S, Hasebe M, Tomita M, Yanagawa H. Systematic identification of cell cycle-dependent yeast nucleocytoplasmic shuttling proteins by prediction of composite motifs. Proc Natl Acad Sci USA. 2009;106(25):10171–6.

39. Camacho C, Coulouris G, Avagyan V, Ma N, Papadopoulos J, Bealer K, et al. BLAST+: architecture and applications. BMC bioinformatics. 2009;10:421.

40. Abramson J, Adler J, Dunger J, Evans R, Green T, Pritzel A, et al. Accurate structure prediction of biomolecular interactions with AlphaFold 3. Nature. 2024;630(8016):493–500.

41. Krissinel E, Henrick K. Inference of macromolecular assemblies from crystalline state. J Mol Biol. 2007;372(3):774–97.

42. Hoad M, Nematollahzadeh S, Petersen GF, Roby JA, Alvisi G, Forwood JK. Structural basis for nuclear import of adeno-associated virus serotype 6 capsid protein. J Virol. 2025;99(1):e0134524.

43. Teh T, Tiganis T, Kobe B. Crystallization of importin alpha, the nuclear-import receptor. Acta crystallographica Section D, Biological crystallography. 1999;55(Pt 2):561–3.

44. Alvisi G, Manaresi E, Cross EM, Hoad M, Akbari N, Pavan S, et al. Importin alpha/beta-dependent nuclear transport of human parvovirus B19 nonstructural protein 1 is essential for viral replication. Antiviral Res. 2023:105588.

45. Sinigalia E, Alvisi G, Mercorelli B, Coen DM, Pari GS, Jans DA, et al. Role of homodimerization of human cytomegalovirus DNA polymerase accessory protein UL44 in origin-dependent DNA replication in cells. J Virol. 2008;82(24):12574–9.

46. Tsujii A, Miyamoto Y, Moriyama T, Tsuchiya Y, Obuse C, Mizuguchi K, et al. Retinoblastoma-binding Protein 4-regulated Classical Nuclear Transport Is Involved in Cellular Senescence. Journal of Biological Chemistry. 2015;290(49):29375–88.

47. Scaturro P, Trist IM, Paul D, Kumar A, Acosta EG, Byrd CM, et al. Characterization of the mode of action of a potent dengue virus capsid inhibitor. J Virol. 2014;88(19):11540–55.

48. Panza E, Marini M, Pecci A, Giacopelli F, Bozzi V, Seri M, et al. Transfection of the mutant MYH9 cDNA reproduces the most typical cellular phenotype of MYH9-related disease in different cell lines. Pathogenetics. 2008;1(1):5.

49. Shaner NC, Lin MZ, McKeown MR, Steinbach PA, Hazelwood KL, Davidson MW, et al. Improving the photostability of bright monomeric orange and red fluorescent proteins. Nature methods. 2008;5(6):545–51.

50. Sarrion-Perdigones A, Gonzalez Y, Venken KJT. Rapid and Efficient Synthetic Assembly of Multiplex Luciferase Reporter Plasmids for the Simultaneous Monitoring of Up to Six Cellular Signaling Pathways. Curr Protoc Mol Biol. 2020;131(1):e121.

51. Roman N, Kirkby B, Marfori M, Kobe B, Forwood JK. Crystallization of the flexible nuclear import receptor importin-beta in the unliganded state. Acta crystallographica Section F, Structural biology and crystallization communications. 2009;65(Pt 6):625–8.

52. Studier FW. Protein production by auto-induction in high density shaking cultures. Protein Expr Purif. 2005;41(1):207–34.

53. Cross EM, Akbari N, Ghassabian H, Hoad M, Pavan S, Ariawan D, et al. A functional and structural comparative analysis of large tumor antigens reveals evolution of different importin alpha-dependent nuclear localization signals. Protein Sci. 2024;33(2):e4876.

54. Cross EM, Marin O, Ariawan D, Aragao D, Cozza G, Di Iorio E, et al. Structural determinants of phosphorylation-dependent nuclear transport of HCMV DNA polymerase processivity factor UL44. FEBS Lett. 2024;598(2):199–209.

55. Aragao D, Aishima J, Cherukuvada H, Clarken R, Clift M, Cowieson NP, et al. MX2: a high-flux undulator microfocus beamline serving both the chemical and macromolecular crystallography communities at the Australian Synchrotron. J Synchrotron Radiat. 2018;25(Pt 3):885–91.

56. Kabsch W. Xds. Acta crystallographica Section D, Biological crystallography. 2010;66(Pt 2):125–32.

57. Evans PR. An introduction to data reduction: space-group determination, scaling and intensity statistics. Acta Crystallogr D. 2011;67:282–92.

58. Collaborative Computational Project N. The CCP4 suite: programs for protein crystallography. Acta crystallographica Section D, Biological crystallography. 1994;50(Pt 5):760–3.

59. McCoy AJ, Grosse-Kunstleve RW, Adams PD, Winn MD, Storoni LC, Read RJ. Phaser crystallographic software. J Appl Crystallogr. 2007;40(Pt 4):658–74.

60. Emsley P, Lohkamp B, Scott WG, Cowtan K. Features and development of Coot. Acta crystallographica Section D, Biological crystallography. 2010;66(Pt 4):486–501.

61. Adams PD, Afonine PV, Bunkoczi G, Chen VB, Davis IW, Echols N, et al. PHENIX: a comprehensive Python-based system for macromolecular structure solution. Acta crystallographica Section D, Biological crystallography. 2010;66(Pt 2):213–21.

62. Messa L, Celegato M, Bertagnin C, Mercorelli B, Alvisi G, Banks L, et al. The Dimeric Form of HPV16 E6 Is Crucial to Drive YAP/TAZ Upregulation through the Targeting of hScrib. Cancers (Basel). 2021;13(16).

63. Di Antonio V, Palu G, Alvisi G. Live-Cell Analysis of Human Cytomegalovirus DNA Polymerase Holoenzyme Assembly by Resonance Energy Transfer Methods. Microorganisms. 2021;9(5).

64. Alvisi G, Jans D, Ripalti A. Human cytomegalovirus (HCMV) DNA polymerase processivity factor ppUL44 dimerizes in the cytosol before translocation to the nucleus. Biochemistry. 2006;45(22):6866–72.

65. Alvisi G, Paolini L, Contarini A, Zambarda C, Di Antonio V, Colosini A, et al. Intersectin goes nuclear: secret life of an endocytic protein. The Biochemical journal. 2018;475(8):1455–72.

66. Smith KM, Di Antonio V, Bellucci L, Thomas DR, Caporuscio F, Ciccarese F, et al. Contribution of the residue at position 4 within classical nuclear localization signals to modulating interaction with importins and nuclear targeting. Biochim Biophys Acta. 2018;1865(8):1114–29.

67. Athukorala A, Donnelly CM, Pavan S, Nematollahzadeh S, Djossou VA, Nath B, et al. Structural and functional characterization of siadenovirus core protein VII nuclear localization demonstrates the existence of multiple nuclear transport pathways. J Gen Virol. 2024;105(1).

68. Trevisan M, Di Antonio V, Radeghieri A, Palu G, Ghildyal R, Alvisi G. Molecular Requirements for Self-Interaction of the Respiratory Syncytial Virus Matrix Protein in Living Mammalian Cells. Viruses. 2018;10(3).

69. Karczewski KJ, Francioli LC, Tiao G, Cummings BB, Alfoldi J, Wang Q, et al. The mutational constraint spectrum quantified from variation in 141,456 humans. Nature. 2020;581(7809):434-43.

70. Gudmundsson S, Singer-Berk M, Watts NA, Phu W, Goodrich JK, Solomonson M, et al. Variant interpretation using population databases: Lessons from gnomAD. Hum Mutat. 2022;43(8):1012–30.

71. Hazawa M, Lin DC, Kobayashi A, Jiang YY, Xu L, Dewi FRP, et al. ROCK-dependent phosphorylation of NUP62 regulates p63 nuclear transport and squamous cell carcinoma proliferation. EMBO reports. 2018;19(1):73–88.

72. Alvisi G, Jans D, Guo J, Pinna L, Ripalti A. A protein kinase CK2 site flanking the nuclear targeting signal enhances nuclear transport of human cytomegalovirus ppUL44. Traffic. 2005;6(11):1002–13.

73. Jakel S, Albig W, Kutay U, Bischoff FR, Schwamborn K, Doenecke D, et al. The importin beta/importin 7 heterodimer is a functional nuclear import receptor for histone H1. EMBO J. 1999;18(9):2411–23.

74. Miyamoto Y, Yamada K, Yoneda Y. Importin alpha: a key molecule in nuclear transport and non-transport functions. J Biochem. 2016;160(2):69–75.

75. Ninpan K, Suptawiwat O, Boonarkart C, Phuangphung P, Sathirareuangchai S, Uiprasertkul M, et al. Expression of importin-alpha isoforms in human nasal mucosa: implication for adaptation of avian influenza A viruses to human host. Virol J. 2016;13:90.

76. Pumroy RA, Cingolani G. Diversification of importin-alpha isoforms in cellular trafficking and disease states. The Biochemical journal. 2015;466(1):13–28.

77. Munasinghe TS, Edwards MR, Tsimbalyuk S, Vogel OA, Smith KM, Stewart M, et al. MERS-CoV ORF4b employs an unusual binding mechanism to target IMPalpha and block innate immunity. Nature communications. 2022;13(1):1604.

78. Hoad M, Roby JA, Forwood JK. Structural basis for nuclear import of bat adeno-associated virus capsid protein. J Gen Virol. 2024;105(3).

79. Klein C, Georges G, Kunkele KP, Huber R, Engh RA, Hansen S. High thermostability and lack of cooperative DNA binding distinguish the p63 core domain from the homologous tumor suppressor p53. J Biol Chem. 2001;276(40):37390–401.

80. Lotz R, Osterburg C, Chaikuad A, Weber S, Akutsu M, Machel AC, et al. Alternative splicing in the DBD linker region of p63 modulates binding to DNA and iASPP in vitro. Cell Death Dis. 2025;16(1):4.

81. Florio TJ, Lokareddy RK, Yeggoni DP, Sankhala RS, Ott CA, Gillilan RE, et al. Differential recognition of canonical NF-kappaB dimers by Importin alpha3. Nature communications. 2022;13(1):1207.

82. Hodel MR, Corbett AH, Hodel AE. Dissection of a nuclear localization signal. J Biol Chem. 2001;276(2):1317–25.

83. Dingwall C, Robbins J, Dilworth SM, Roberts B, Richardson WD. The nucleoplasmin nuclear location sequence is larger and more complex than that of SV-40 large T antigen. The Journal of cell biology. 1988;107(3):841–9.

84. Nematollahzadeh S, Athukorala A, Donnelly CM, Pavan S, Atelie-Djossou V, Di Iorio E, et al. Mechanistic Insights Into an Ancient Adenovirus Precursor Protein VII Show Multiple Nuclear Import Receptor Pathways. Traffic. 2024;25(9):e12953.

85. Wang L, Li M, Cai M, Xing J, Wang S, Zheng C. A PY-nuclear localization signal is required for nuclear accumulation of HCMV UL79 protein. Med Microbiol Immunol. 2012;201(3):381–7.

86. Ping X, Cheng Y, Bao J, Shi K, Zou J, Shentu X. KPNA4 is involved in cataract formation via the nuclear import of p53. Gene. 2021;786:145621.

87. Nie L, Sasaki M, Maki CG. Regulation of p53 nuclear export through sequential changes in conformation and ubiquitination. J Biol Chem. 2007;282(19):14616–25.

88. Liang SH, Hong D, Clarke MF. Cooperation of a single lysine mutation and a C-terminal domain in the cytoplasmic sequestration of the p53 protein. J Biol Chem. 1998;273(31):19817–21.

89. Matsuura Y. Structural and biochemical characterization of the recognition of the 53BP1 nuclear localization signal by importin-alpha. Biochem Biophys Res Commun. 2019;510(2):236–41.

90. Eschenfeldt WH, Lucy S, Millard CS, Joachimiak A, Mark ID. A family of LIC vectors for high-throughput cloning and purification of proteins. Methods Mol Biol. 2009;498:105–15.

