## Supplementary Figurs S1-S5 and SUpplementary Tables S1-S6 for "A bipartite, mutation-tolerant NLS regulates interaction of ΔNp63α with importin alpha, nuclear transport and transcriptional activity"

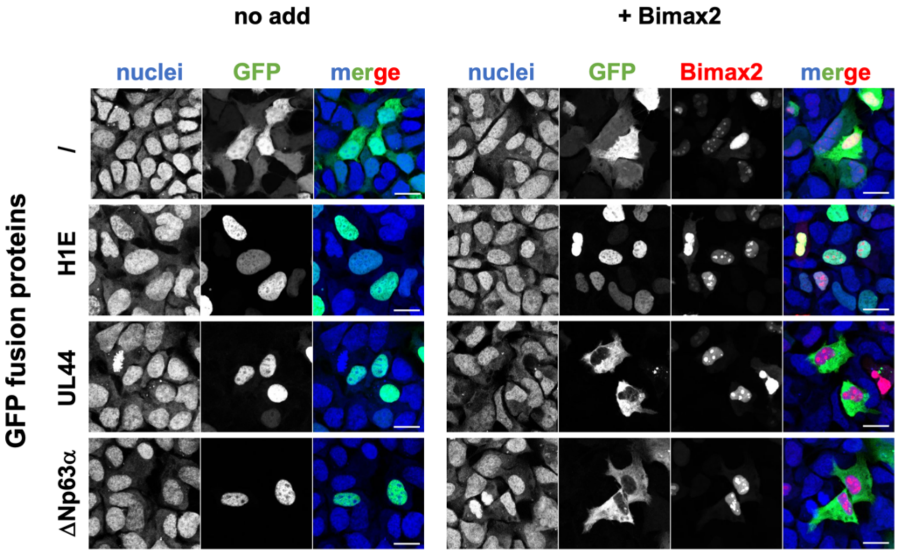


**<u>Figure S1.</u> ΔNp63α is transported into the nucleus by the IMPα/β1 heterodimer.** Separate channel images relative to micrographs shown in Figure 1.

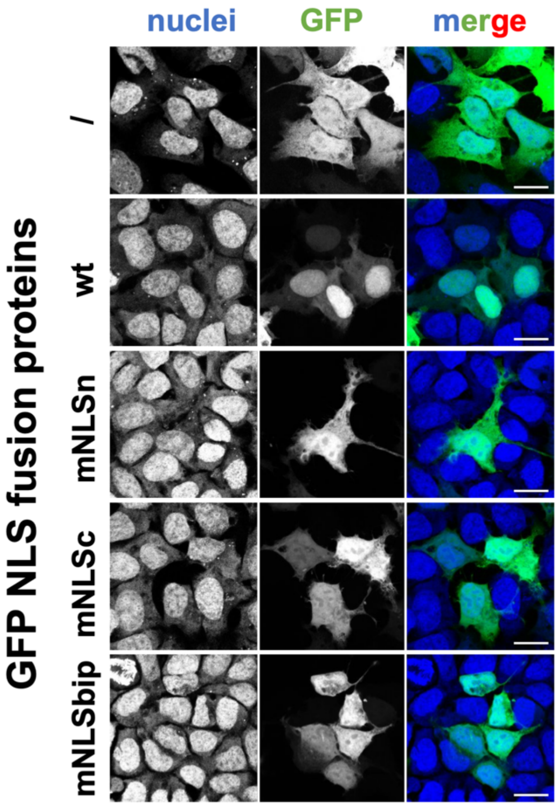


**Supplementary Figure S2.** **ΔNp63α residues 278-302 represent a functional bipartite NLS.** Separate channel images relative to micrographs shown in Figure 2.

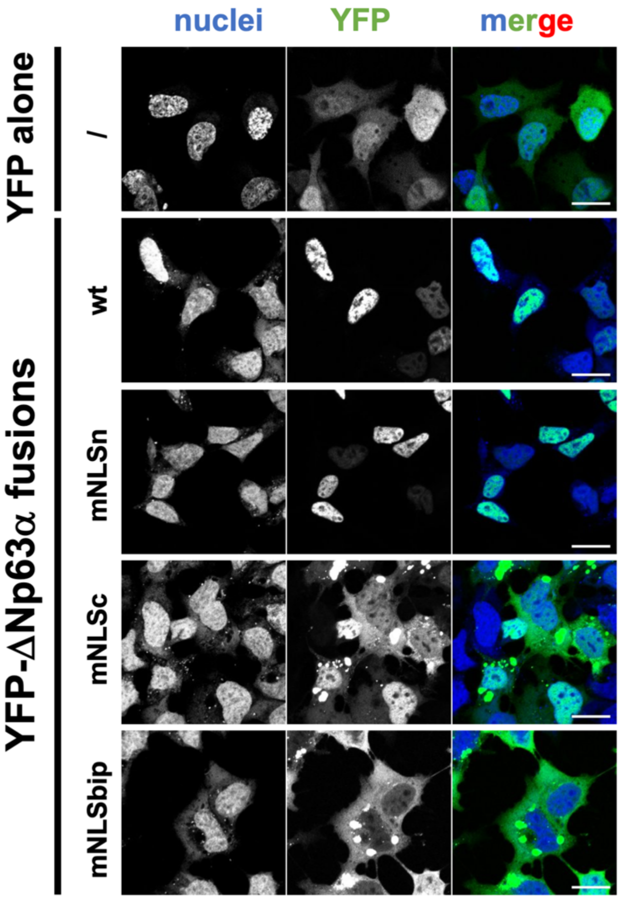


**<u>Supplementary Figure S3</u>. Substitution of basic residues within ΔNp63α bipartite NLS results in failure to accumulate in the cell nuclei.** Separate channel images relative to micrographs shown in Figure 6.

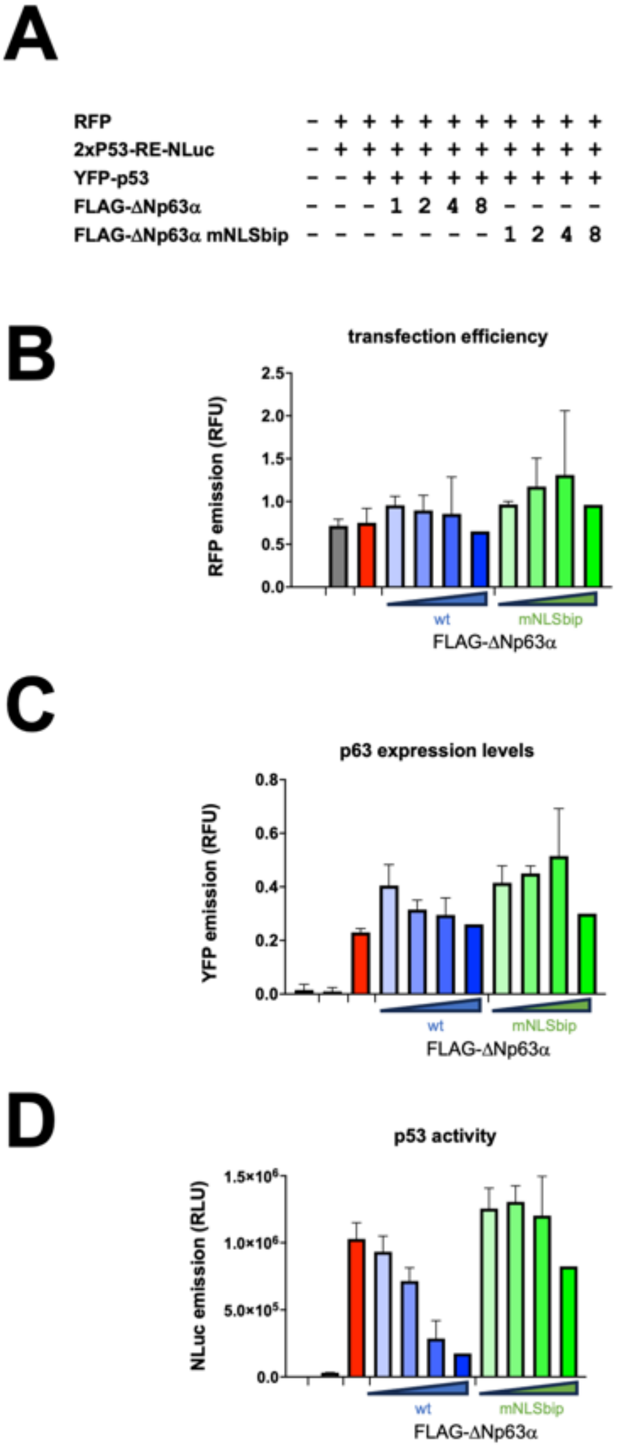


<u>**Supplementary Figure S4**</u>**. Nuclear localization is essential for ΔNp63α transcriptional inhibition.** Raw data relative to Figure 7. (A) H1299 were transfected with plasmids mediating expression of Tag-RFP under the control of the constitutive Immediate early (IE) promoter (CMV), to monitor transfection efficiency; nanoluciferase (NLuc) under the control of two p53 response elements (2xTP53 RE), to monitor p53-dependent transcription; YFP-p53 under the control of CMV IE promoter to transactivate 2xTP53 RE; and increasing amounts of either FLAG-ΔNp63α or FLAG-ΔNp63α;mNLSbip to repress the 2xTP53 RE. Twenty four hours later, cells were processed as described in the Materials and Methods to perform luciferase assays and monitor 2xTP53 RE transactivation. (B) The expression of RFP under each condition is shown, as the mean RFP fluorescence expressed in relative fluorescent units (RFU). (C) The

expression of YFP-p53 under each condition is shown, as the mean YFP fluorescence emission expressed in relative fluorescent units (RFU). (D) The activity of YFP-p53 under each condition is shown as the mean NLuc activity expressed in relative luminescent units (RFU). Data are shown as means + standard deviation of the mean relative to three independent experiments

**A**

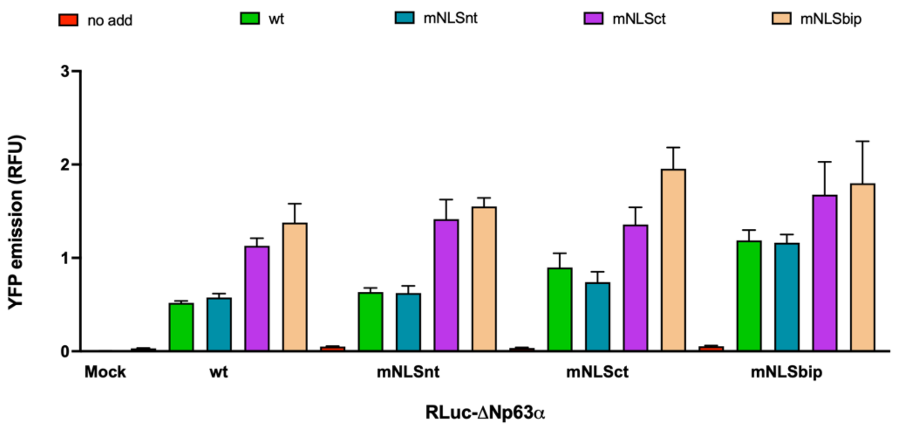


**B**

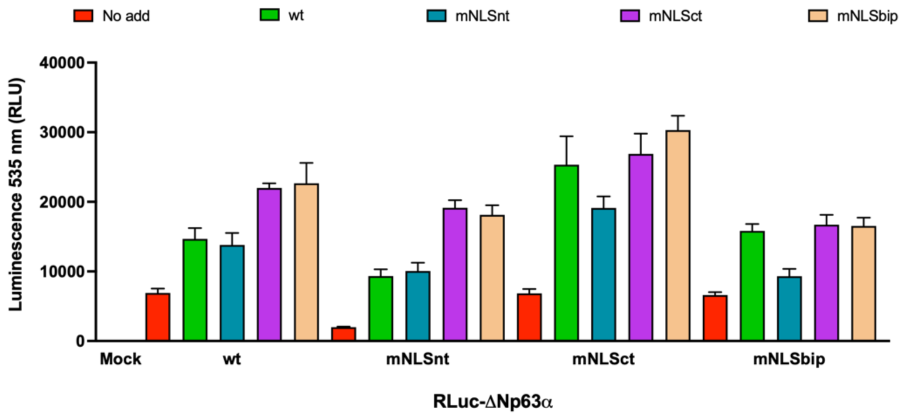


**C**

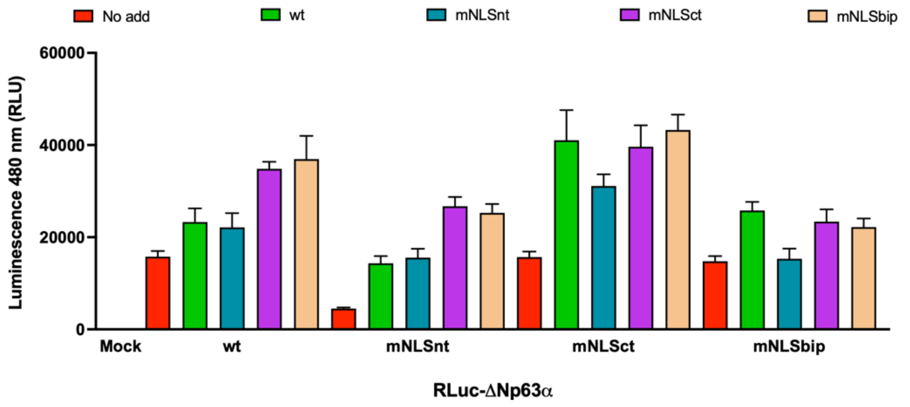


**Supplementary Figure S5. Nuclear localization is not required for ΔNp63α self-association.** Additional measurements relative to Figure 7. HEK293T cells were transfected with 45 ng of the indicated RLuc-ΔNp63α fusion proteins in the absence (no add), or in the presence of 250 ng of plasmids mediating the expression of the indicated YFP-ΔNp63α fusion proteins. Twenty-four hours later, cells were processed as described in the Materials and Methods section to perform endpoint BRET assays, thanks to measurement of YFP fluorescence (A) as well as luminescence with a 480+20 nm filter (B) and with a 535+20 nm (C). Data shown are means and standard error of the mean relative to 3 independent experiments performed in triplicate.

| ***Plasmid name*** | **Details** |
|---|---|
| pcDNA3.1-NT-GFP-TOPO | Mammalian expression of cycle 3 GFP. Used as a negative control for nuclear accumulation (44). |
| pcDNA3.1-NT-GFP-TOPO-SV40-NLS | Mammalian expression of cycle 3 GFP fused to SV40 Large T antigen NLS (PKKKRKV-132). Used as a positive control for nuclear accumulation (44). |
| pEPI-GFP-UL44 | Mammalian expression of HCMV DNA polymerase processivity factor UL44 with a N-terminal GFP tag. Used as a positive control for IMPα/β1-dependent cargo (45). |
| pCDNA3.1-mCherry-Bimax2 | Mammalian expression of the IMPα/β1 inhibitor Bimax2 with an N-terminal mCherry tag. Used to test dependency on IMPα/β1-mediated nuclear import (46). |
| pEGFP-C1-p53 | Mammalian expression of p53 with a N-terminal GFP tag, flanked by attB sites. Used to generate pDNR207-p53. |
| pEGFP-C1-ΔNp63α | Mammalian expression of ΔNp63α with a N-terminal GFP tag, flanked by attB sites. Used to generate pDNR207-ΔNp63α and to test the dependency of ΔNp63α nuclear import on IMPα/β1. |
| pEGFP-C1-ΔNp63α NLSbip | Mammalian expression of ΔNp63α NLSbip (278-DGTKRPFRQNTHGIQMTSIKKRRSP-302) with a N-terminal GFP tag. Used to test nuclear targeting activity of ΔNp63α NLSbip. |
| pEGFP-C1-ΔNp63α NLSbip;mNLSn | Mammalian expression of ΔNp63α NLSbip containing Ala substitutions in NLSn (278-DGTAAPFRQNTHGIQMTSIKKRRSP-302) with a N-terminal GFP tag. Used to test nuclear targeting activity of ΔNp63α NLSbip. |
| pEGFP-C1-ΔNp63α NLSbip;mNLSc | Mammalian expression of ΔNp63α NLSbip containing Ala substitutions in NLSc (278-DGTKRPFRQNTHGIQMTSIAAAASP-302) with a N-terminal GFP tag. Used to test nuclear targeting activity of ΔNp63α NLSbip. |
| pEGFP-C1-ΔNp63α;mNLSbip | Mammalian expression of ΔNp63α NLSbip containing Ala substitutions in NLSbip (278-DGTAAPFRQNTHGIQMTSIAAAASP-302), with a N-terminal GFP tag. Used to test nuclear targeting activity of ΔNp63α NLSbip. |
| pDNR207-ΔNp63α | Gateway ENTRY vector coding ΔNp63α sequence, followed by a STOP codon and flanked by attB sites. Used to perform LR reactions. |
| pDNR207-ΔNp63α;mNLSn | Gateway ENTRY vector coding ΔNp63α sequence containing Ala substitutions in NLSn (278-DGTAAPFRQNTHGIQMTSIKKRRSP-302), followed by a STOP codon and flanked by attB sites. Used to perform LR reactions. |
| pDNR207-ΔNp63α;mNLSc | Gateway ENTRY vector coding ΔNp63α sequence containing Ala substitutions in NLSc (278-DGTKRPFRQNTHGIQMTSIAAAASP-302), followed by a STOP codon and flanked by attB sites. Used to perform LR reactions. |

| | |
|---|---|
| pDNR207-ΔNp63α;mNLSbip | Gateway ENTRY vector coding ΔNp63α sequence containing Ala substitutions in NLSbip (278-DGTAAPFRQNTHGIQMTSIAAAASP-302), followed by a STOP codon and flanked by attB sites. Used to perform LR reactions. |
| pDESTntYFP-ΔNp63α | Mammalian expression of ΔNp63α with a N-terminal YFP tag. Used for subcellular localization and BRET assays. |
| pDESTntYFP-ΔNp63α;mNLSn | Mammalian expression of ΔNp63α containing Ala substitutions in NLSn (278-DGTAAPFRQNTHGIQMTSIKKRRSP-302), with a N-terminal YFP tag. Used for subcellular localization. |
| pDESTntYFP-ΔNp63α;mNLSc | Mammalian expression of ΔNp63α containing Ala substitutions in NLSc (278-DGTKRPFRQNTHGIQMTSIAAAASP-302), with a N-terminal YFP tag. Used for subcellular localization. |
| pDESTntYFP-ΔNp63α;mNLSbip | Mammalian expression of ΔNp63α containing Ala substitutions in NLSbip (278-DGTAAPFRQNTHGIQMTSIAAAASP-302), with a N-terminal YFP tag. Used for subcellular localization. |
| pDESTntRLuc-ΔNp63α | Mammalian expression of ΔNp63α with a N-terminal RLuc tag. Used for BRET assays. |
| pDESTntRLuc-ΔNp63α;mNLSbip | Mammalian expression of ΔNp63α containing Ala substitutions in NLSbip (278-DGTAAPFRQNTHGIQMTSIAAAASP-302), with a N-terminal RLuc tag. Used for BRET assays. |
| pDESTntFLAG-ΔNp63α | Mammalian expression of ΔNp63α with a N-terminal FLAG tag. Used for p53 transcriptional inhibition assays. |
| pDESTntFLAG-ΔNp63α;mNLSbip | Mammalian expression of ΔNp63α containing Ala substitutions in NLSbip (278-DGTAAPFRQNTHGIQMTSIAAAASP-302), with a N-terminal FLAG tag. Used for p53 transcriptional inhibition assays. |
| *pDESTntRFP*-ΔNp63α | Mammalian expression of ΔNp63α with a N-terminal RFP tag. Used for ΔNp63α p63 co-transport assays. |
| TagRFP-N | Mammalian expression of TagRPT-T under the control of the CMV IE promoter. Used for p53 transcriptional inhibition assays (49). |
| 2xP53_RE::NLuc | Mammalian expression of NLuc under the control of 2 x P53 RE. Used for p53 transcriptional inhibition assays (50). |
| pDESTntYFP-p53 | Mammalian expression of p53 with a N-terminal YFP tag. Used for p53 transcriptional inhibition assays. |
| pET30a-hIMPα1ΔIBB | Bacterial expression of recombinant His tagged human IMPα1ΔIBB, with a Tobacco Etch Virus (TEV) protease site between the coding sequence and the His tag (43), |
| pET30a-mIMPα2ΔIBB | Bacterial expression of recombinant His tagged moue IMPα2ΔIBB, with a Tobacco Etch Virus (TEV) protease site between the coding sequence and the His tag (43). |
| pET30a-hIMPα3ΔIBB | Bacterial expression of recombinant His tagged human IMPα3ΔIBB, with a Tobacco Etch Virus (TEV) protease site between the coding sequence and the His tag (43). |
| pET30a-hIMPα5ΔIBB | Bacterial expression of recombinant His tagged human IMPα5ΔIBB, with a Tobacco Etch Virus (TEV) protease site between the coding sequence and the His tag (43). |

| | |
|---|---|
| pET30a-hIMP$\alpha$7$\Delta$IBB | Bacterial expression of recombinant His tagged human IMP$\alpha$7$\Delta$IBB, with a Tobacco Etch Virus (TEV) protease site between the coding sequence and the His tag (43). |
| pMCSG21-hIMP$\beta$1 | Bacterial expression of recombinant His tagged human IMP$\beta$1, with a Tobacco Etch Virus (TEV) protease site between the coding sequence and the His tag (90). |

Supplementary Table S1. Plasmids used in this study.

| ***Name*** | **Sequence** |
| --- | --- |
| *ΔNp63αNLS* | FITC-Ahx-GTKRPFRQNTHGIQMTSIKKRRSP |
| *ΔNp63α NLS;mNLSn* | FITC-Ahx-GTAAPFRQNTHGIQMTSIKKRRSP |
| *ΔNp63α NLS;mNLSc* | FITC-Ahx-GTKRPFRQNTHGIQMTSIAAAASP |
| *ΔNp63α NLS;mNLSbip* | FITC-Ahx-GTAAPFRQNTHGIQMTSIAAAASP |

Supplementary Table S2. Sequences of peptides used in this study.

| IMPs | Kd (nM) | Bmax (mP) |
|---|---|---|
| IMPα1△IBB | 4.3 ± 0.1 | 184.3 ± 0.6 |
| IMPα2△IBB | 57.4 ± 2.8 | 148.9 ± 1.3 |
| IMPα3△IBB | 6.7 ± 0.3 | 201.5 ± 1.3 |
| IMPα5△IBB | 9.1 ± 0.5 | 171.6 ± 1.3 |
| IMPα7△IBB | 4.6 ± 0.4 | 169.1 ±1 .9 |
| IMPβ1 | > 20,000 | 166.2 ± 36.7 |

Supplementary Table S3. Summary statistics of FP assays shown in Figure 3. Data are shown as means ± standard error of the mean relative to three independent experiments.

| **ΔNp63α NLS:IMPα2ΔIBB complex (PDB Code: 9N54)** | |
|---|---|
| *Data Collection (high resolution statistics in parentheses)* | |
| Wavelength | 0.95374 |
| Data-collection temperature (K) | 298 |
| Detector Type | Dectris EIGER X 16M |
| Detector | Pixel |
| Resolution range (Å) | 43.75 - 2.20 |
| Space group | P21 21 21 |
| Unit cell (Å); (o) | 78 90 100; 90 90 90 |
| Total reflections | 346552 |
| Unique reflections | 36733 (3134) |
| Multiplicity | 9.4 (9.6) |
| Completeness (%) | 100.0 (100.0) |
| Mean I/σ (I) | 19.5 (4.8) |
| Wilson B-factor Å2 | 34.91 |
| Rpim | 0.022 (0.195) |
| *Refinement* | |
| Rwork | 0.184 |
| Rfree | 0.208 |
| No. of non-hydrogen atoms | 6952 |
| Macromolecules | 2 |
| Solvent | 135 |
| Protein residues | 441 |
| Bond length r.m.s.d (Å) | 0.004 |
| Bond angle r.m.s.d (o) | 0.591 |
| Ramachandran favored (%) | 97.7 |
| Ramachandran allowed (%) | 2.3 |
| Ramachandran outliers (%) | 0.0 |

Supplementary Table S4. Data collection and refinement statistics for structure of ΔNp63α NLS:mIMPα2ΔIBB complex.

| ΔNp63α NLS:IMPα2ΔIBB complex (PDB Code: 9N54) | | |
|---|---|---|
| **Hydrogen Bonds** | | |
| *Structure 1* | *Dist. [Å]* | *Structure 2* |
| A:LYS 281[HZ2] | 2.35 | E:THR 328 [OG1] |
| A:LYS 281[HZ3] | 2.14 | E:THR 328 [OG1] |
| A:LYS 281[HZ1] | 2.17 | E:ASN 361 [O] |
| A:LYS 281[HZ3] | 2.19 | E:VAL 321 [O] |
| A:ARG 282 [H] | 1.97 | E:ASN 361 [OD1] |
| A:ARG 282 [HH21] | 2.15 | E:SER 360 [OG] |
| A:ARG 282 [HH22] | 1.75 | E:GLU 396 [OE1] |
| A:LYS 297 [HZ3] | 2.13 | E:THR 155 [OG1] |
| A:LYS 297 [HZ1] | 2.13 | E:ASP 192 [OD1] |
| A:LYS 298 [H] | 2.02 | E:ASN 188 [OD1] |
| A:ARG 299 [HH12] | 1.96 | E:ARG 106 [O] |
| A:ARG 299 [HH11] | 2.14 | E:LEU 104 [O] |
| A:ARG 299 [HH22] | 2.18 | E:ARG 106 [O] |
| A:ARG 300 [H] | 2.12 | E:ASN 146 [OD1] |
| A:ARG 300 [HH11] | 1.71 | E:GLN 181 [OE1] |
| A:ARG 282 [O] | 2.35 | E:ASN 361 [HD21] |
| A:ILE 296 [O] | 2.46 | E:ASN 235 [HD21] |
| A:LYS 298 [O] | 2.07 | E:ASN 188 [HD21] |
| A:ARG 300 [O] | 2.03 | E:ASN 146 [HD21] |
| **Salt Bridges** | | |
| A:ARG 282 [NH1] | 3.35 | E:GLU 396 [OE1] |
| A:ARG 282 [NH1] | 3.66 | E:GLU 396 [OE2] |
| A:ARG 282 [NH2] | 2.60 | E:GLU 396 [OE1] |
| A:LYS 297 [NZ] | 2.96 | E:ASP 192 [OD1] |

Supplementary Table S5. Hydrogen bonds and salt bridges of ΔNp63α NLS:mIMPα2ΔIBB complex.
